# A Systematic Review and Meta-Analysis of Biological Sex Differences in Sleep Spindles and Slow Wave Activity in Adults with and without Insomnia

**DOI:** 10.64898/2026.01.19.700353

**Authors:** Nyissa A. Walsh, Emma-Maria Phillips, Arsenio Páez, Nathan E. Cross, Thien Thanh Dang-Vu, Aurore A. Perrault

## Abstract

Mounting evidence shows sex-based differences in sleep experiences and outcomes, including the prevalence of insomnia disorder. However, the impact of biological sex on brain oscillations during sleep remains poorly understood, especially in the context of insomnia disorder. This is a notable gap, given that neurophysiological aspects of sleep are associated with brain health and overall sleep quality. We systematically reviewed and meta-analysed data from studies reporting spindle and slow wave activity in adults with and without insomnia disorder. We conducted systematic searches in PubMed, Embase, Scopus, and PsycInfo. Risk of bias was evaluated using the Cochrane Risk of Bias tool.

Forty-three studies met our inclusion criteria, with thirteen studies of normal sleepers (N= 668) reporting sufficient data for random-effects meta-analyses. Compared with males, female normal sleepers had higher spindle density, sigma and delta power. Most studies recruited individuals with primary insomnia, and data pooling for insomnia and mixed groups was not possible due to insufficient statistical reporting. Moreover, group-by-sex interactions were limited, inconsistent, and varied across studies and sample characteristics. Further research is needed to explore sex-specific differences in sleep microarchitecture and their role in normal sleep and the manifestation of insomnia disorder.

## INTRODUCTION

Chronic insomnia or insomnia disorder is the most prevalent sleep disorder among adults affecting approximately 10-17% of the population worldwide[1,2]. It is defined as self-reported difficulties falling and/or staying asleep (insomnia symptoms), occurring more than three nights per week, persisting for at least three months, despite adequate opportunity to sleep, and negative impacting daytime functioning such as mood disturbances and cognitive performance difficulties[3,4]. The prevalence of chronic insomnia is one and a half times higher in females compared to males, and females often report poorer sleep quality than males in the general population[1,5]. The incidence of insomnia disorder differs by sex and age, with middle and older-aged females at the highest risk for developing insomnia, compared to younger females and males of all ages[1,4] Given the difficulty in establishing the direction of causality, the current definition of insomnia disorder does not distinguish between insomnia with potentially contributing comorbidities and insomnia without (formerly called primary insomnia[6]).

In females, chronic insomnia is frequently reported during major reproductive life events marked by significant hormonal and physical changes, such as puberty, pregnancy, and perimenopause[7]. Indeed, prior to the first menses (menarche), there is no difference between young females and males in the likelihood of developing insomnia, likely due to less hormonal fluctuations between sexes[8]. However, the risk of developing insomnia increases almost threefold in females after puberty and the establishment of menstrual cycles, while males risk remains relatively stable[8,9]. Similarly, during the menopausal transition (perimenopause), there is a large increase in the risk of developing insomnia symptoms, from 35% in pre-menopausal females to 53% in post-menopausal females[9]. Menopausal transition is associated with decreases in levels of both progesterone and estrogen, which are female reproductive hormones that have, among other functions, neurological protective effects. They increase synaptogenesis in brain regions important for memory and directly affect the brain regions associated with sleep[10–12]. Thus, from adolescence to late life, fluctuations in gonadal steroids (such as estrogen and progesterone) may be a contributing factor to the risk of developing insomnia disorder in females by influencing sleep physiology.

While insomnia diagnosis is solely based on subjective complaints, polysomnographic (PSG) recordings can reveal objective markers of sleep disturbances (e.g., longer sleep latency, more frequent awakenings) in adults with chronic insomnia[13]. In normal sleepers, sex-based differences can already be observed at the macro-architecture level, as it has been shown that females tend to go to bed earlier, fall asleep quicker and spend more time in bed. Compared to males, females also sleep longer and spend less time awake (i.e., greater sleep efficiency [SE]; total sleep time/total time in bed X 100%)[14–16]. Those differences are apparent in young healthy adults but also in middle-aged and older adults. Although sleep duration decreases with age regardless of sex, males tend to exhibit a somewhat significant linear decrease in total sleep time across the lifespan. In contrast, the sleep duration of females stays fairly stable until around age 50, then declines— around menopause. By age 70, males and females have similar sleep durations[17]. Interestingly, although females tend to objectively sleep better than their male age matched counterparts across the lifespan (e.g., longer NREM3 duration from midlife onward) [18,19], they often report poorer sleep quality in the general population[20,21]. This suggests that sex-based differences in insomnia may be influenced not only by variations in sleep architecture, but also by psychosocial factors such as stress reactivity, hyperarousal, mood disturbances, and caregiving responsibilities, which may uniquely affect females experience of sleep disturbance[7].

While differences in sleep architecture and psychosocial factors are informative clinically, the investigation of brain oscillations offers a deeper understanding of the brain mechanisms underlying sleep depth, quality and sleep-dependent memory processes[22,23]. For instance, slow wave activity (SWA), a marker of homeostatic sleep drive (i.e., the pressure to sleep that builds during wakefulness and dissipates with sleep) reflects restorative NREM sleep and is linked to synaptic plasticity and memory consolidation [24]. SWA is composed of slow waves (SW; also called delta waves) oscillating between 1 and 4 Hz as well as high-voltage biphasic waves called slow oscillations (SO; <1.25Hz, > 75μV)[25–27]. It is important to study both SW and SO activity as they originate from distinct neural mechanisms[25]. Indeed, SOs typically emerge from the prefrontal cortex, while SW stem from the thalamus and various cortical regions[28]. Some persons with insomnia disorder show reduced or suppressed SWA, possibly due to the inability to fully disengage from a waking brain state, which may contribute to the feeling of non-restorative sleep[29]. Whether studied as a waveform (i.e., event detection of SO <1.25Hz and slow waves <4Hz) or generalised activity over a longer period event (i.e., spectral power in the SO or delta frequency bands), SWA has been often associated with cognitive and physiological functions such energy saving, metabolite clearance but also hormonal modulatory action[30].

Another crucial brain oscillation important for neural plasticity and restorative sleep are sleep spindles[31,32]. Oscillating within the sigma range (11-16 Hz) between 0.5 to 3 s, spindles are abundant during NREM2 and NREM3[33]. Sleep spindles are often categorized based on spatiotemporal characteristics; fast spindles (∼14-16 Hz) occur mainly in central and partial cortex, whereas slow spindles (∼11-13 Hz) are located more frontally. Both spindles are key for understanding insomnia because their density, amplitude, and synchrony support stable, restorative sleep and protect against stress-related disruptions[31,34,35]. Higher spindle activity correlates with deeper sleep and better perceived rest[35], while lower spindle density—especially early in the night—predicts greater vulnerability to stress and environmental disturbances, both of which can worsen insomnia symptoms[36]. However, studies on spindle activity in individuals with insomnia disorder are inconsistent; some find fewer or weaker spindles, while others find no differences compared to good sleepers[37]. Despite these inconsistencies, spindles may still be useful for predicting who will respond well to cognitive-behavioral therapy for insomnia (CBTi)[38], because they are thought to regulate sleep stability by filtering external stimuli[36].

### Evidence Gap

Given the high prevalence of insomnia in females, this study fills a critical gap in sleep research as the first systematic synthesis and meta-analysis to investigate sex-based differences in spindle and SWA extracted from EEG recordings across both adults with insomnia and normal sleeping populations. To address this critical evidence gap, we aimed to determine whether measures of spindle activity (including spindle event and sigma power) and SWA (including SO, SW, delta power) differ between biological sex (i.e., female, male) in adults with and without insomnia. While prior reviews have focused on macro-sleep architecture or neural oscillation alterations in insomnia without stratifying by sex, our work is the first synthesis to integrate sex and neural oscillations in insomnia, to explain why females tend to report poorer sleep quality and higher insomnia risk, despite often having better objective sleep than males.

Conducting a systematic review of the current evidence will help pinpoint priority areas for future research and enhance evidence-based treatment of insomnia disorder.

## METHODS

The systematic review was conducted following the recommendations of the Cochrane Handbook for Systematic Reviews of Interventions[39]. Results are reported in accordance to the Preferred Reporting Items for Systematic Reviews and Meta-Analyses (PRISMA) guidelines[40]. The protocol was registered on the NIHR-PROSPERO prospective register of systematic reviews (https://www.crd.york.ac.uk/PROSPERO/view/CRD42022355969).

### Eligibility Criteria

**P**opulation: Human adults over 18 years old, with or without insomnia disorder. Papers including younger individuals were eligible if data extraction was possible for the target age group, or if the mean age was 18 years old or greater.

**E**xposure and **C**omparison: Studies including **both** males and females.

**O**utcomes: Eligible papers had to report continuous data for spindle measures (i.e., spindle density, amplitude, frequency, peak frequency, count, duration, laterality, distribution, diffuseness, as well as absolute, relative, and normalized sigma power) and/or SWA measures (i.e., SO and SW density, SW amplitude, frequency, phase duration, slope, incidence, distribution, laterality, and diffuseness as well as absolute, relative, and normalized delta power and SO power) recorded during sleep studies.

**S**tudy design: Only peer-reviewed, cross-sectional, case-control, cohort, or randomized-controlled trail (RCT) studies were eligible for the current review. Pharmacological/psychological intervention studies were included if they reported baseline (pre-treatment) information about participants meeting our eligibility criteria. To be eligible for the meta-analyses, papers included in the systematic review had to report sex-specific data on brain oscillations in normal sleepers and/or persons with insomnia.

### Exclusion Criteria

Meeting abstracts, posters, case reports, editorial, commentary, and narrative review papers were excluded. Papers not reporting sex-based analyses of sleep spindle or SWA we deemed ineligible. Studies with participants under the age of 18 or those including only males or only females were excluded as cross-study sex comparisons risk conflating biological differences with methodological artifacts (e.g., measurement differences in spindle or SWA). Given the effects of chronic diseases and associated treatments on sleep, we excluded studies on participants with pre-defined major medical disorders (e.g., cancer, chronic pain; major neurological disorders such as– epilepsy, dementia, traumatic brain injury; major psychiatric disorders such as– schizophrenia, major depression, bipolar disorder)[4]. Studies assessing treatment effects on PSG/EEG outcomes were excluded, but baseline data were retained. Given both positive and negative effects of treatments/interventions on sleep, mood, and memory, including such cases may have confounded the results of this review[41]. Therefore, post-treatment outcomes on participants undergoing interventions or treatments were not included. For example, benzodiazepines, commonly prescribed sleep-inducing drugs has been shown to alter sleep spindles during NREM[42,43]. Sleep medications were also an exclusion criterion. Only papers containing broad spectrum EEG variables (e.g., delta and sigma power) and NREM events (i.e., SO, SW and spindle density, duration, amplitude, frequency, phase duration, slope, incidence, distribution laterality, and diffuseness) were included. Therefore, studies that only examined sleep architecture variables (e.g., total sleep time, duration of sleep stages) or extracted spectral activity within EEG frequency bands not of interest (e.g., alpha) were removed. Studies indicating “slow wave sleep” were initially included for further investigation to ensure they reported data for SWA either as an event- or spectral power-based rather than NREM3 stage percentage, which is often reported as SWA within sleep literature. If only the percentage of time spent in NREM3 was reported, the study was excluded.

For the meta-analysis, if eligible studies did not report summary statistics (e.g., means, SDs), we attempted to contact authors to obtain the missing data. If data remained unavailable, studies were excluded from meta-analysis but included in the systematic review.

### Information Sources

Systematic searches were conducted on PubMed, Embase, Scopus and PsycInfo from inception to November 2025. To ensure literature saturation, we also undertook citation searching of the reference lists of two relevant reviews that were excluded from our systematic review—specifically, a meta-analysis on sleep microarchitecture in insomnia (Baglioni et al., 2014[44]) and a systematic review on sex-based differences in the sleep-wake cycle (Carrier et al., 2017[16]) —to identify any additional eligible studies not captured by our primary search.

### Search Strategy

Searches were undertaken with Medical Subject Headings (MeSH) terms, and keywords related to brain oscillations in persons with and without insomnia (See **Figure S1**). Keywords for biological sex analyses were not included to avoid missing papers reporting sex-based differences but not describing them in their methods. Papers written in any language were eligible without publication date restriction.

### Selection Process

After the removal of duplicate results using the Zotero Software (Version 6.0.37), the following selection process was applied: First, search results were screened against our eligibility criteria by title and abstract. Papers meeting those criteria, or papers for which we needed more information to make a determination, were then selected for full-text screening. Authors were contacted to obtain additional information regarding the study’s method to assess eligibility when necessary. The first stage of the selection process was conducted by two reviewers (NAW, EMP). Agreement between study eligibility was based on a consensus between three authors (NAW, EMP, AAP) and any disagreements were resolved by consensus from all six authors.

### Data Extraction

Data extraction was undertaken and checked by the first reviewer (NAW) and second investigator (EMP). Using Zotero Software and customised Microsoft Excel sheets created following PRISMA guidelines. Study characteristics included the author name, publication year, study location, population demographics (e.g., frequency of biological sex, average age, age range, sample size), study design, the method used to measure brain oscillations (e.g., number of nights, sampling rate, electrodes used, epochs time length for scoring, stages used for scoring [NREM2, NREM3 or both]), the target variables of interest (e.g., spindle density, SO amplitude), and the main outcomes corresponding to the aims of the systematic review.

Given the frequent interchangeability of the terms sex and gender in the insomnia literature, we used the terms women, females, men, and males to categorize participants based on biological sex. Since none of the included studies examined the societal influences of gender on sleep, our analyses were grounded in biological sex rather than gender as a social construct.

Given heterogeneity in terms for brain oscillations found in the sleep literature (i.e., non-systematic use of similar terms to define spindle, sigma, SO, delta and SWA), any outcomes (e.g., density, amplitude, peak frequency, duration) referring to an event detection were either categorised as spindles (detected within sigma range), slow waves (SW; detected within 0.25-4Hz) or SO (detected below 1.25Hz). Meanwhile, if power spectral analyses were used, they were categorised as sigma (∼11-16Hz), delta (∼0.25- 4Hz), or SO (<1.25Hz) spectral power. When appropriate, sleep spindles were reported as fast (∼14-16 Hz) or slow spindles (∼11-13 Hz) according to papers methods. Less common terms were also extracted such as *spindle distribution*[45,46] (i.e., the spatial pattern of sleep spindles across the scalp, categorized by regions [frontal, central, parietal], providing a detailed mapping of spindle activity), *spindle laterality*[46] (i.e., using a measure of spindle amplitude differences between hemispheres) and *spindle diffuseness*[45] (i.e., the number of channels out of six where spindles are detected simultaneously and the spatial distribution of spindles across frontal, central, and parietal regions, providing a detailed scalp-wide mapping of activity).

We extracted sex effect estimates for spindle and SWA outcomes (i.e., spindle density, sigma power and delta power) as mean differences with variance measures. Outcomes of the studies were categorized as normal sleeper, insomnia, or both, for analysis and reporting. If key data were missing or unclear, study authors were contacted when possible; otherwise, studies with insufficient data were excluded. All relevant results for each outcome were sought.

### Risk of Bias Assessment

Data extraction and risk of bias assessment were independently conducted by two reviewers (N.A.W and A.P.) using the Newcastle-Ottawa Scale (NOS)[47] for cross-sectional studies, case-control studies, cohort studies, and the PEDro scale for randomized controlled trials[48]. Any discrepancies in risk of bias assessments were resolved through consensus between the authors. Studies deemed to be at high risk of bias were excluded from meta-analysis. The quality of included studies informed the synthesis and interpretation of our results.

### Sleep spindle and SWA measures

We extracted means, standard deviations, and sample sizes from studies reporting EEG/PSG data on sleep spindle and SWA measures.

### Data Analysis

Descriptive statistics were used to quantify and describe the results of our systematic review literature searches, screening, and study characteristics (e.g., outcomes). We expected clinical diversity and heterogeneity between studies resulting from variable samples (e.g. mean ages, proportion of men and women), study locations and settings. Studies with small sample sizes and large standard errors may also bias the results of pooled effect estimates in meta-analysis [49]. Therefore, following the guidance of the Cochrane Handbook for Systematic Reviews of Interventions, we conducted inverse-variance weighted random-effects (Der Simonian and Laird[50]) meta-analyses with Review Manager 5.41 software.

Individual Hedges *g* effect sizes/standardized mean differences between males and females in spindle and SWA were calculated in RevMan 5.41 with continuous outcome data from each paper eligible for meta-analysis to account for differences in scales or processes between papers[49].

Pooled data from meta-analyses on sleep spindle density and SWA was displayed graphically using Forest plots.

Heterogeneity was assessed with *t*^2^, χ^2^ test (significance level: 0.1) and *I*^2^ statistic, as follows[39]:

1. 0% to 30%: might not be important.
2. 30% to 50%: may represent moderate heterogeneity.
3. 50% to 90%: may represent substantial heterogeneity.
4. 75% to 100%: considerable heterogeneity.

Statistical heterogeneity among reviewed studies was also evaluated visually and displayed graphically by Forest plot[39,51]. Heterogeneity was explored by sensitivity analyses, removing individual papers from meta-analysis to see their effect on heterogeneity when it was high (*I*^2^ >50%)[39,53].

### Publication Bias

Publication bias was explored visually using effect size (Hedges g) by standard error funnel plots [39]. We did not conduct additional statistical tests (e.g., Egger’s test, trim-and-fill) or assess other types of reporting bias (e.g., selective outcome reporting). In meta-analyses with a small sample of studies (< 10 papers) per outcome, the power of these tests is too low to distinguish real asymmetry from chance[39]. As a result, Egger’s tests for publication bias were not undertaken.

## RESULTS

### Study Selection

Our systematic searches yielded 13,290 papers from PubMed, Embase, Scopus, and PsycInfo (**Figure 1)**. After removing 3,782 duplicates, 9,506 papers were reviewed by title and abstract. From these, 9,064 papers were excluded. Four hundred eighty-six papers were deemed eligible or likely eligible and screened by full-text, of which 224 were excluded after full screening. Citation searching among relevant but ineligible papers[44],[16], added four eligible papers resulting in 257 papers meeting our inclusion criteria. Of these, 17% (43 papers) analyzed sex-based differences in spindle or SWA: 33 on normal sleepers (NS) only, two on insomnia (IN) only, and eight on both populations (**Tables 1, 2 and 3**). Attrition and acknowledgement of sex-based difference per population study (NS, IN, mixed) can be found in the *Supplemental Material*.

**Figure 1.**
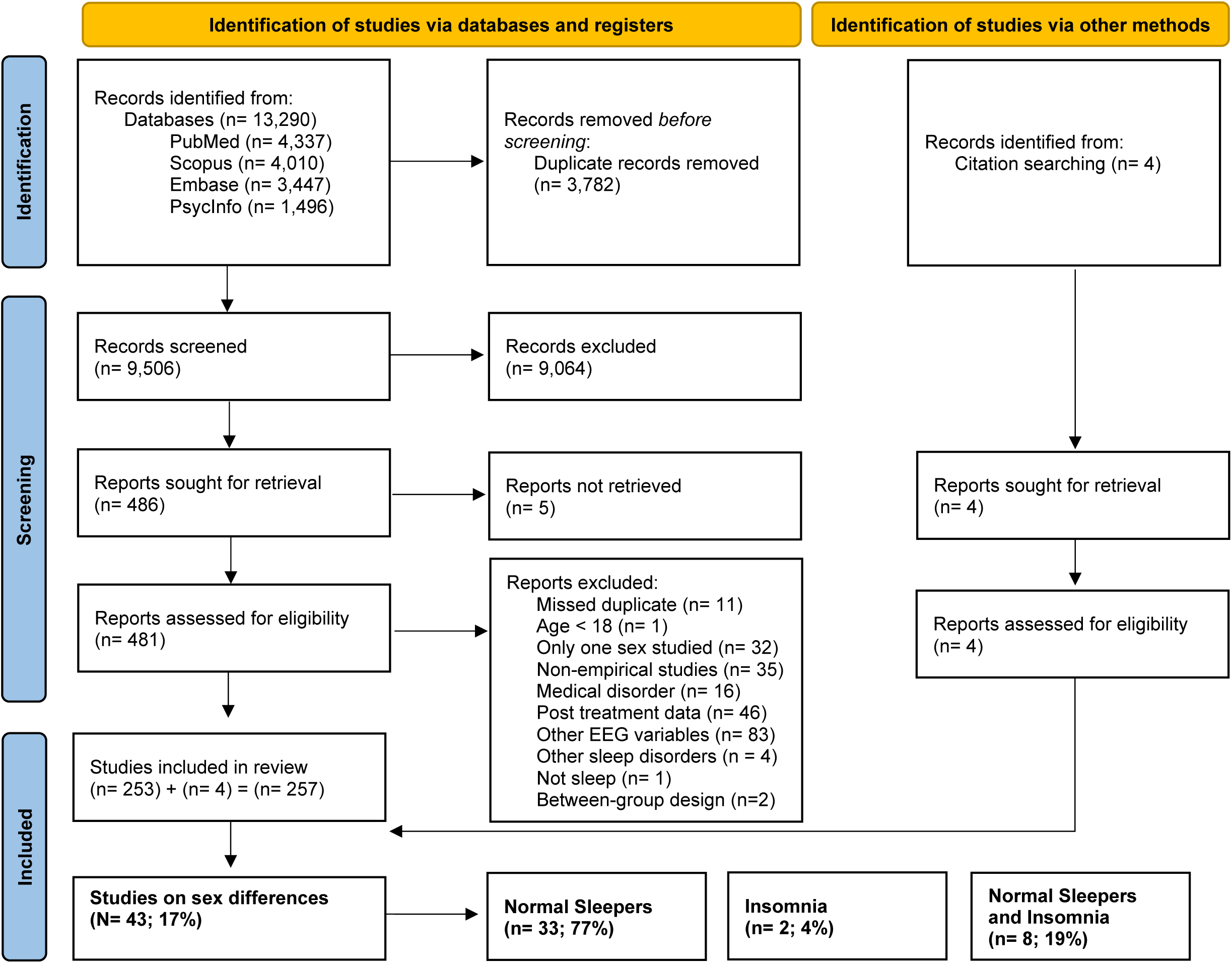
Flow Diagram for Systematic Review and Meta-Analysis

**Table 1.**
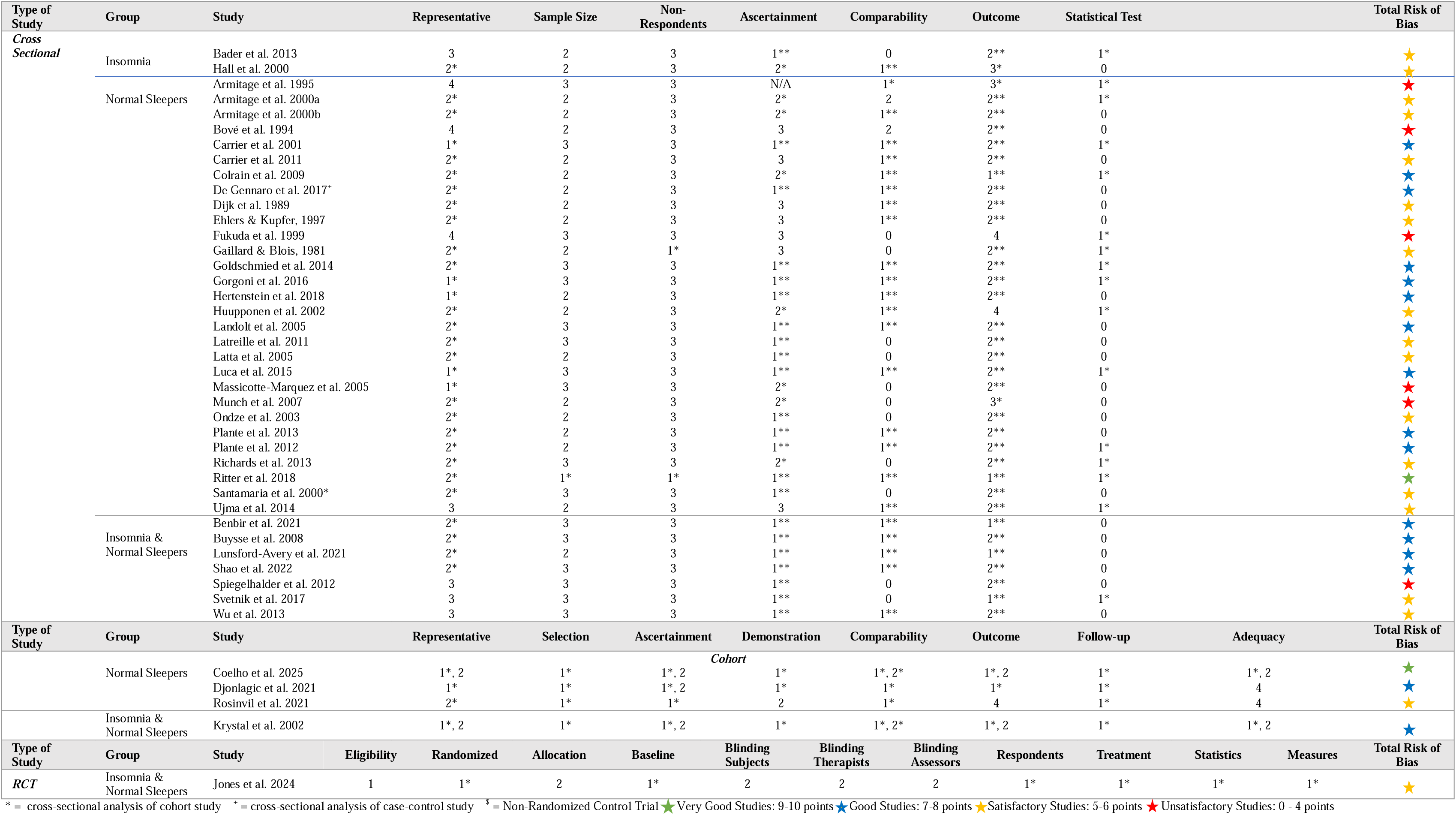
Risk of Bias Summary Plot According to the Newcastle-Ottawa Scale for Cross Sectional, Cohort, Case-Control Studies, and Randomized Controlled Trial Studies

**Table 2.**
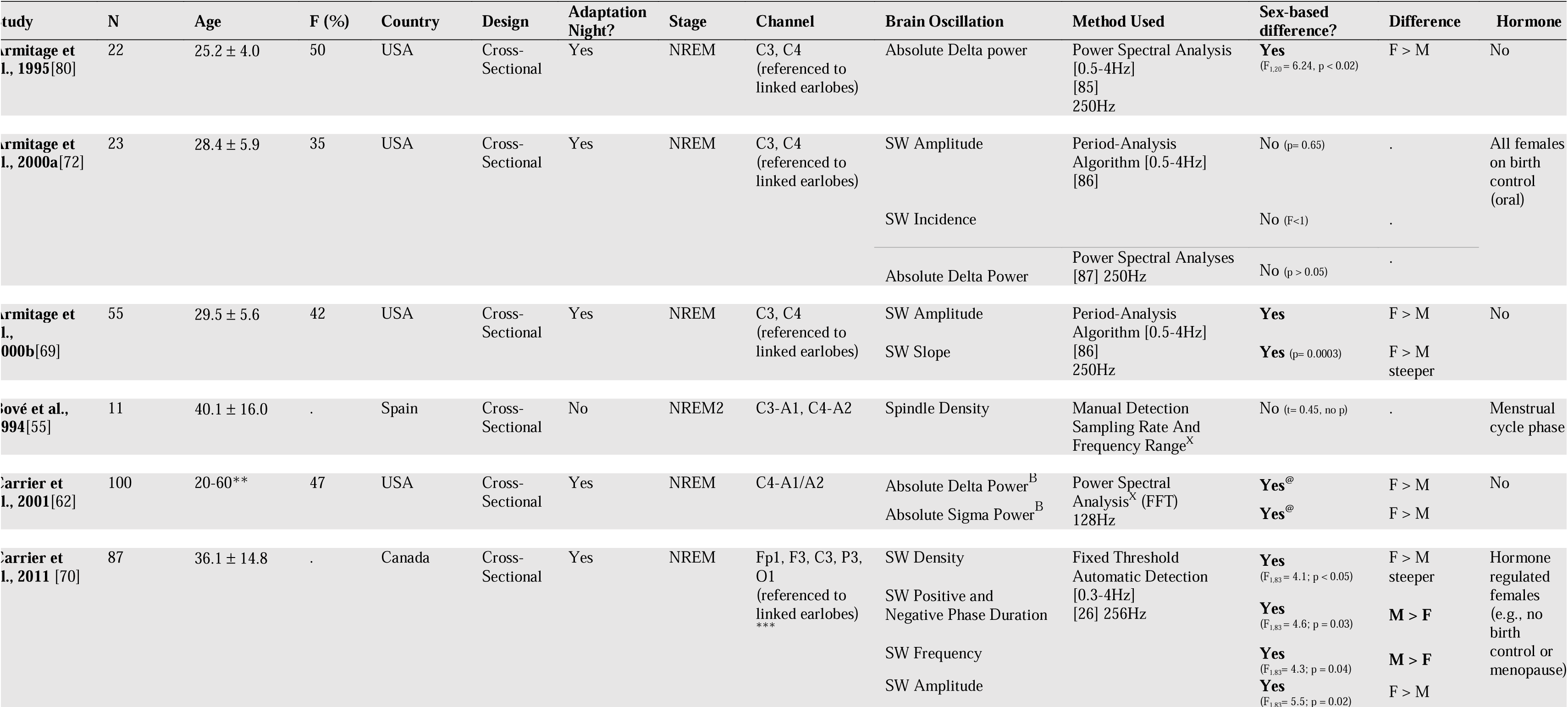
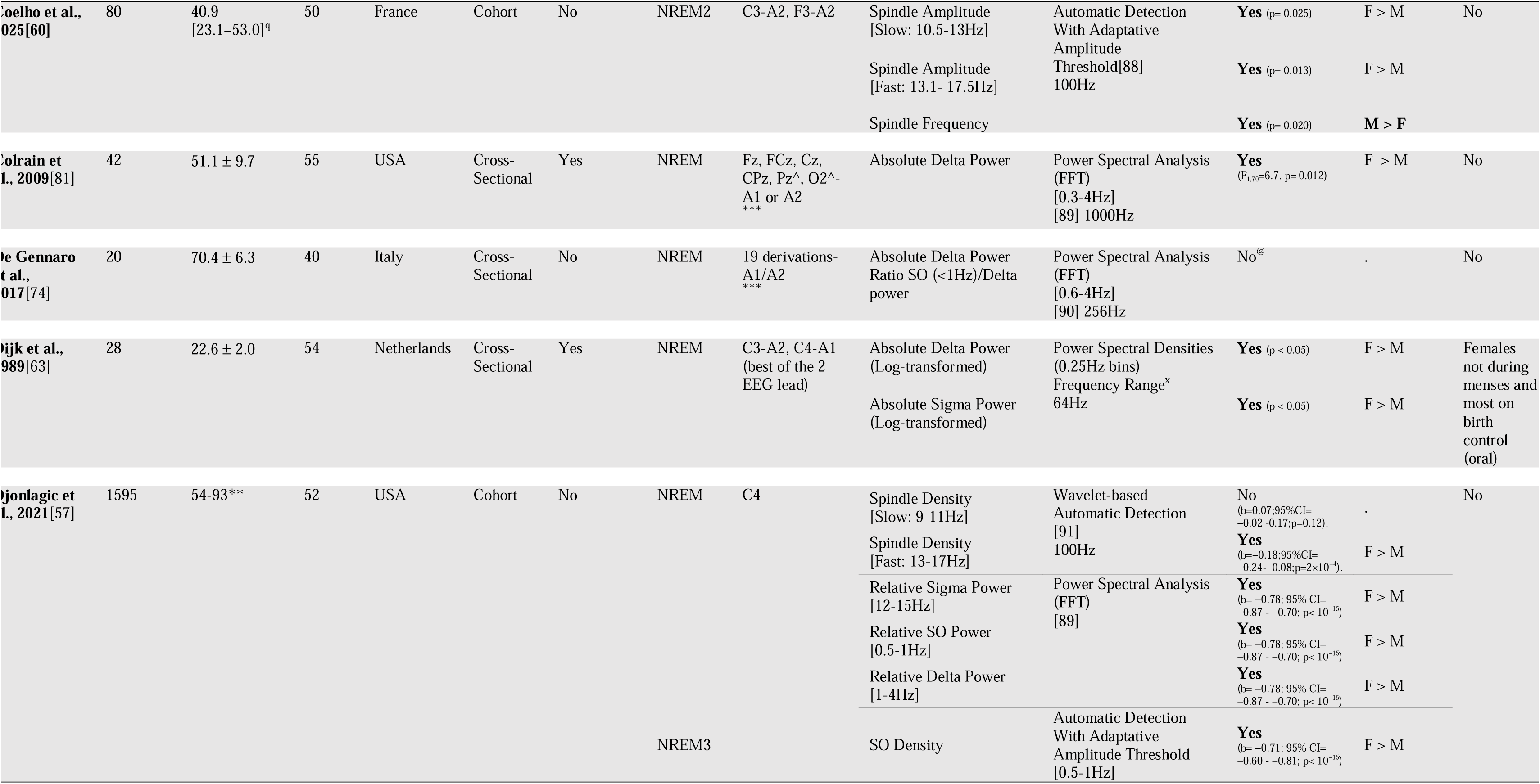
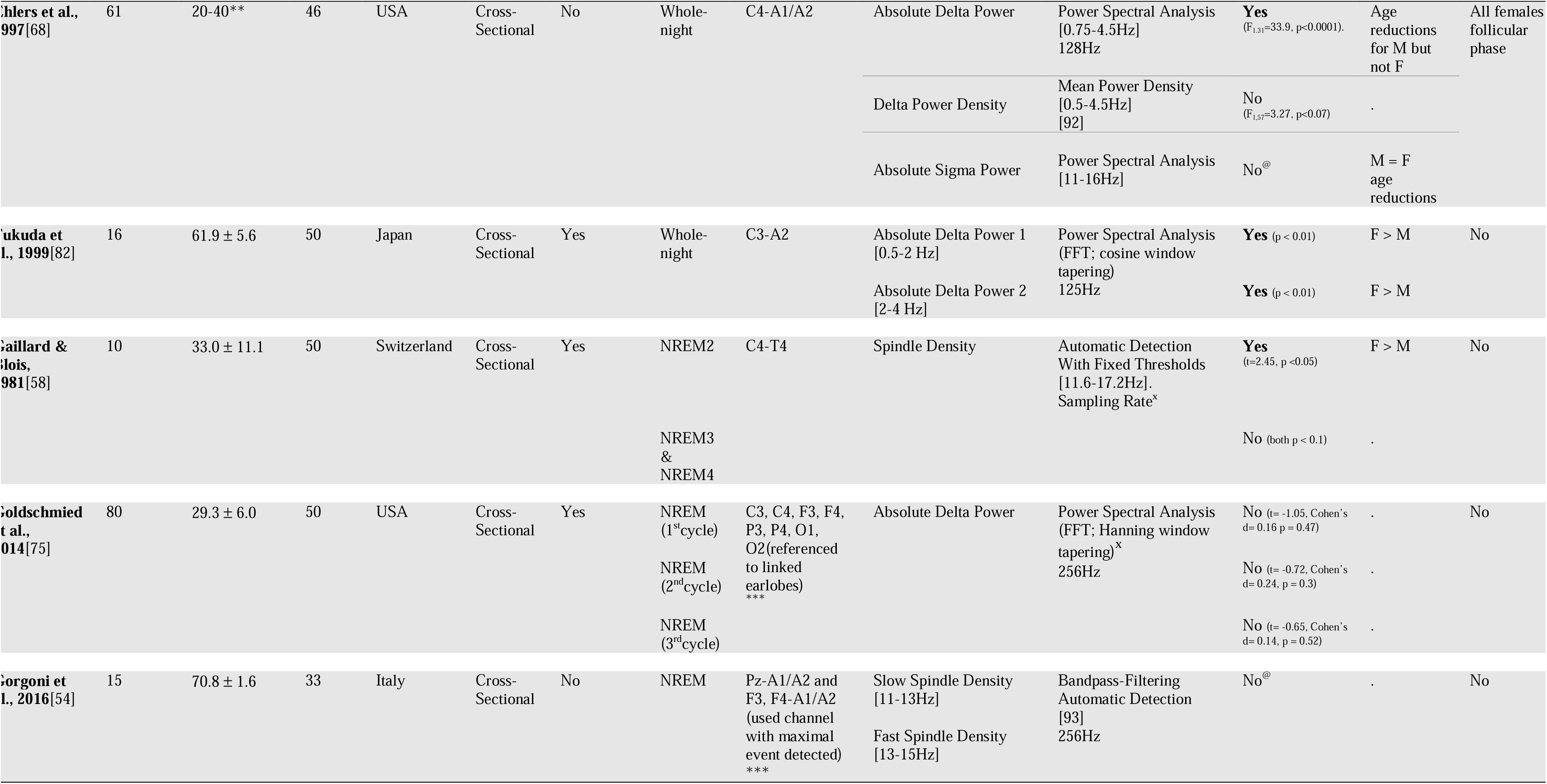
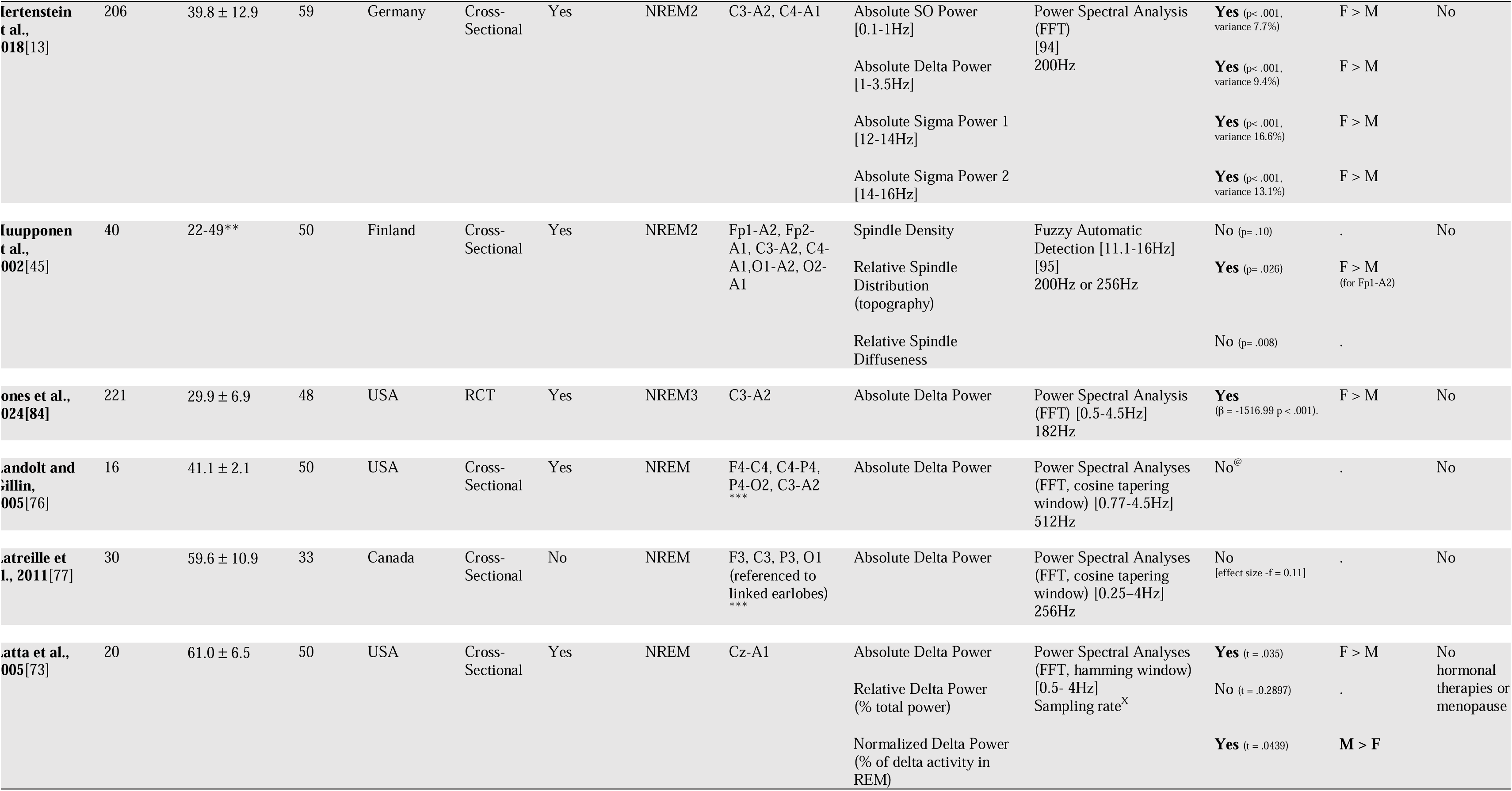
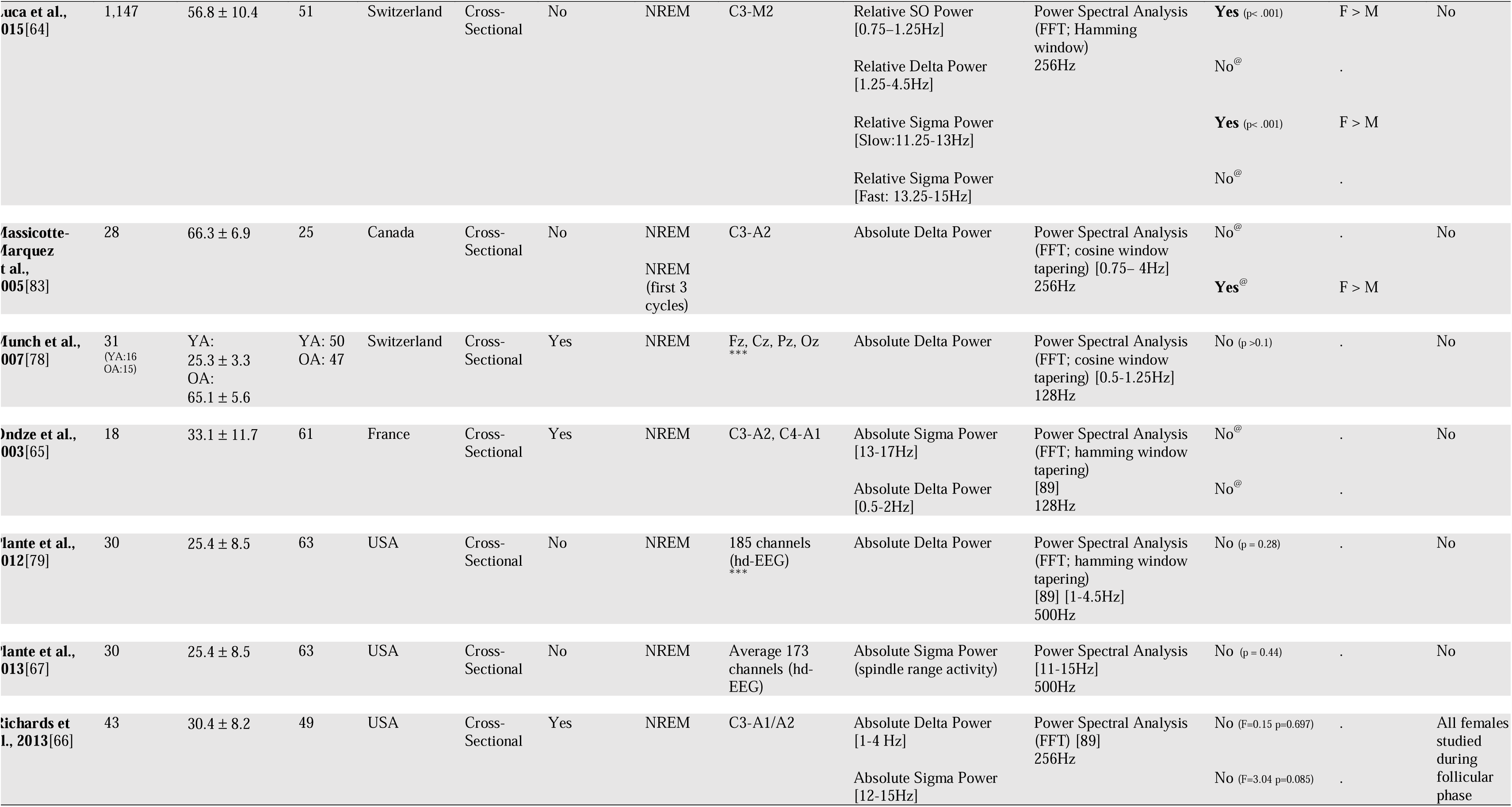
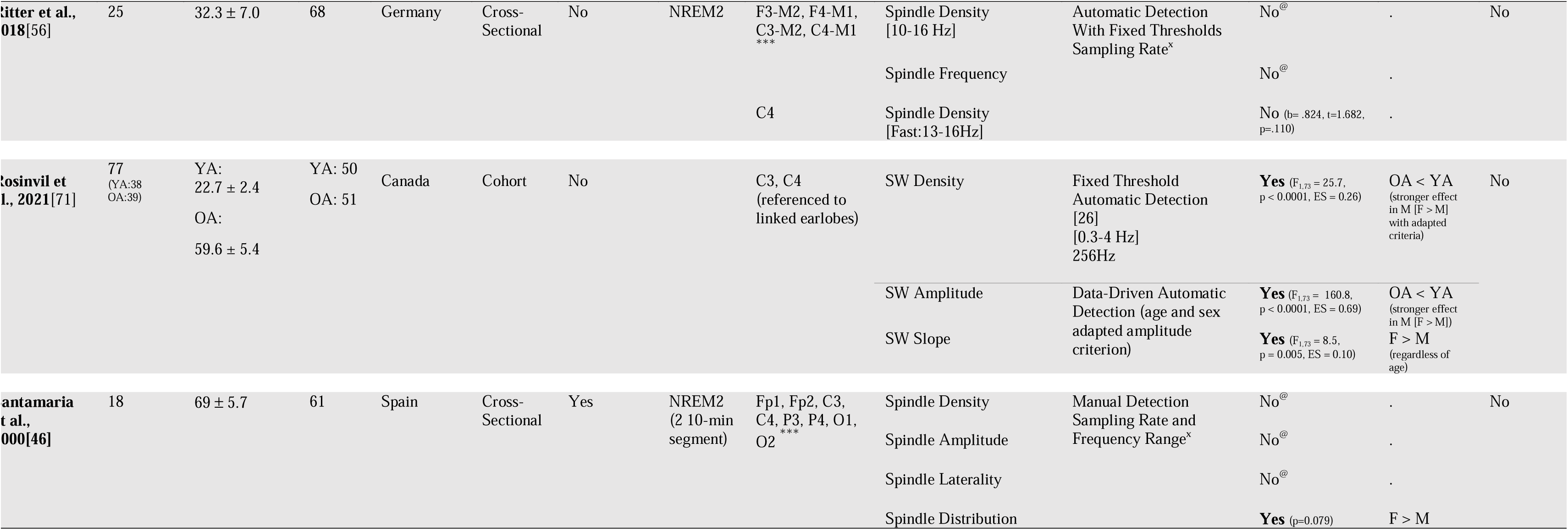
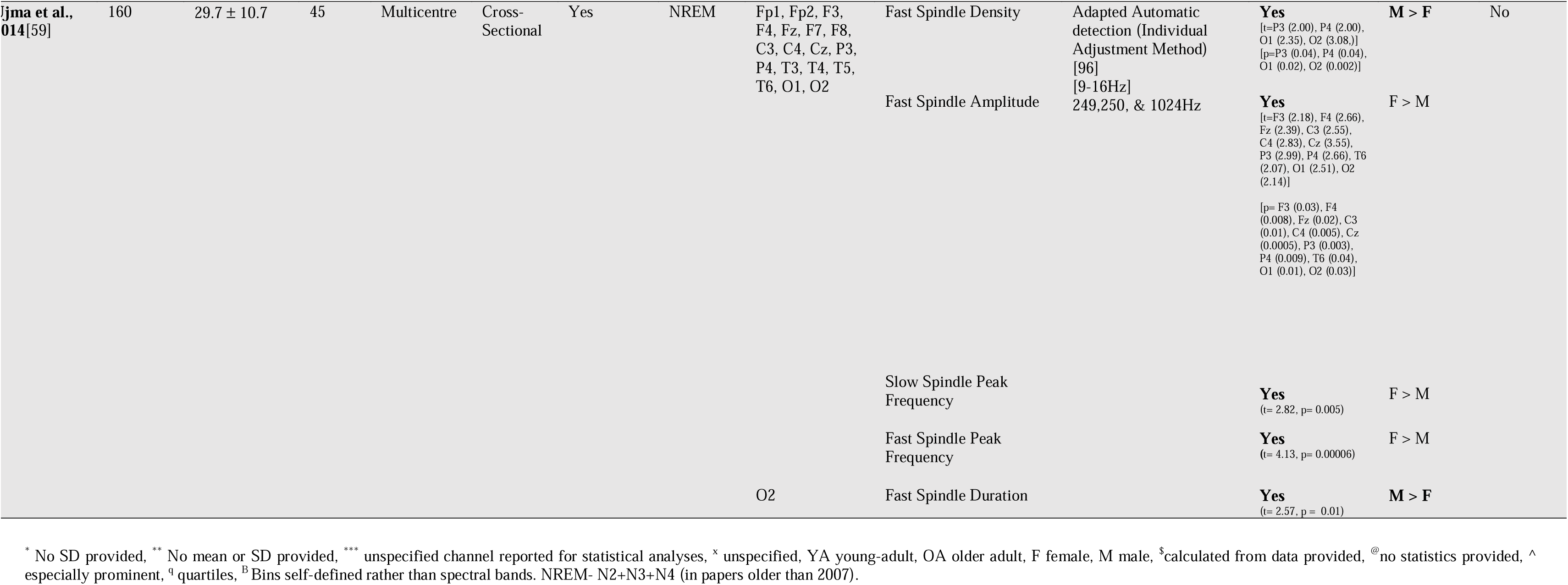
Study Characteristics of Normal Sleepers

**Table 3.**
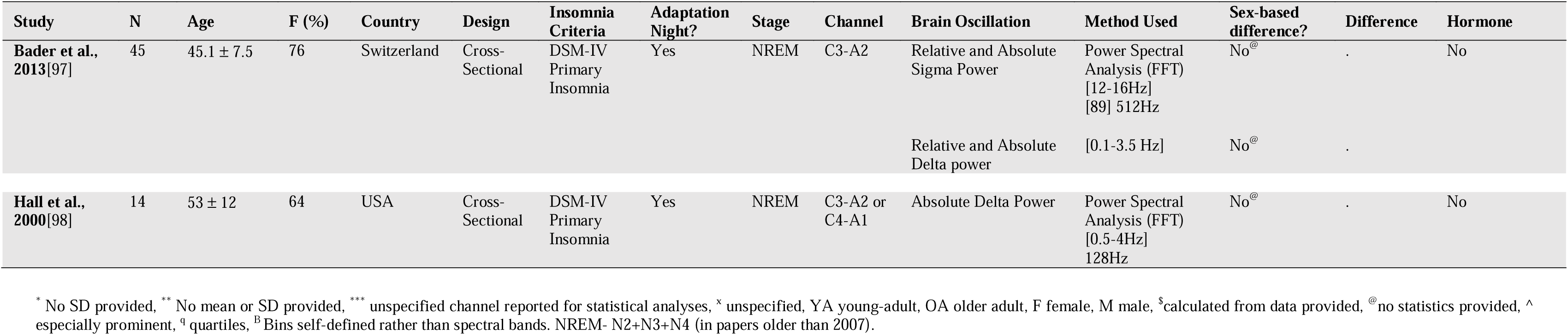
Study Characteristics of Insomnia

| Study | N | Age | F (%) | Country | Design | Insomnia Criteria | Adaptation Night? | Stage | Channel | Brain Oscillation | Method Used | Sex-based difference? | Difference | Hormone |
| --- | --- | --- | --- | --- | --- | --- | --- | --- | --- | --- | --- | --- | --- | --- |
| **Bader et al., 2013**[97] | 45 | 45.1 ± 7.5 | 76 | Switzerland | Cross-Sectional | DSM-IV Primary Insomnia | Yes | NREM | C3-A2 | Relative and Absolute Sigma Power | Power Spectral Analysis (FFT) [12-16Hz] [89] 512Hz | No<sup>@</sup> | . | No |
|  |  |  |  |  |  |  |  |  |  | Relative and Absolute Delta power | [0.1-3.5 Hz] | No<sup>@</sup> | . |  |
| **Hall et al., 2000**[98] | 14 | 53 ± 12 | 64 | USA | Cross-Sectional | DSM-IV Primary Insomnia | Yes | NREM | C3-A2 or C4-A1 | Absolute Delta Power | Power Spectral Analysis (FFT) [0.5-4Hz] 128Hz | No<sup>@</sup> | . | No |

\* No SD provided, \*\* No mean or SD provided, \*\*\* unspecified channel reported for statistical analyses, <sup>x</sup> unspecified, YA young-adult, OA older adult, F female, M male, <sup>s</sup>calculated from data provided, <sup>@</sup>no statistics provided, <sup>^</sup> especially prominent, <sup>q</sup> quartiles, <sup>B</sup> Bins self-defined rather than spectral bands. NREM- N2+N3+N4 (in papers older than 2007).

### Risk of Bias Assessment

Among the 43 studies reviewed, most exhibited low to moderate risk of bias with satisfactory to good quality. Fourteen percent showing high-risk concerns according to the Newcastle-Ottawa Scale (See **Table 1**). The primary risk of bias resulted from inadequate sample size justification, missing data on non-respondents, and the absence of effect size measures (e.g., confidence intervals, p-values). These issues suggest that many studies were potentially underpowered, affecting the reliability of their findings. One study assessed with the PEDro Scale had fair-to-good study quality, with risk of bias arising from lack of reporting about participant or examiner blinding or allocation concealment[52].

### Findings From the Systematic Review

#### Normal Sleepers

Thirty-three of the included papers directly examined sex-based differences in brain oscillations during sleep (see **Table 2**). Out of the 33 studies evaluating sex-based differences, 17 reported on spindle activity, 23 reported on SWA characteristics, and three reported specifically on SO characteristics. It is important to note that while all the papers included in our review examined sex-based differences for at least one type of brain oscillation, they did not do so systematically across all oscillatory measures. For example, some studies assessed sex-based differences in sigma power but not in delta power. Nevertheless, these studies were included in both sections of our review to evaluate the effects of insomnia on each brain oscillation, regardless of whether sex-based differences were specifically analysed for every measure.

##### Normal Sleepers: Spindle Activity

Seventeen research papers explored sex-based differences in spindle characteristics in NS, representing a total of 3,587 participants (mean age= 41.4 ± 18.6, median group size= 35, range= [10, 1595]; see **Table 2**) and nine spindle characteristic outcomes (sigma power, spindle density, spindle frequency, spindle peak frequency, spindle amplitude, spindle duration, spindle laterality, spindle distribution, and spindle diffuseness).

##### Spindle (event-based detection)

The most commonly reported domains were spindle density (k = 8). Spindle detection methods varied between papers (**Table 2**). Out of eight papers specifically reporting on spindle density, 38% (k= 3) reported significant sex-based differences, while 62% (k = 5) did not[45,46,54–56]. Of the three papers finding differences between sexes in spindle density: 67%(k= 2) found higher density in females[57,58] (one[22] specific to fast spindles, and one specifically only during NREM2[58]), and 33% (k= 1) found higher density in males[59]. The majority of papers not finding sex-based differences in spindle density (60%; k= 3) only analysed spindles during NREM2[45,46,55].

Other spindle characteristic outcomes were less commonly reported across studies. Two articles [56,60] reported on spindle frequency (Hz), one found no sex-based differences[56], while the other indicated males had higher frequency than females[60]. Three studies reported on spindle amplitude; one (50%)[46] reported no sex-based differences, while the others found females had higher spindle amplitudes, specifically in the fast spindle range (F3) [59,60]. One of the two found the same for the slow spindle range (C3)[60]. The same study was the sole paper that looked at spindles duration[59], finding that males had greater fast spindle duration (13-15 [12.5-15.38]Hz) compared to females in Oz. Two studies on spindle distribution (100%) reported higher distribution in females[45,46]. One specifically found this relationship to be true in the left frontal area but not central or occipital[45]. There was no sex-based differences in spindle laterality[46] and spindle diffuseness[61] (k= 1).

##### Sigma Spectral Power

Beyond spindle activity based on event detection, some studies studied spindle activity using power spectral analysis referred as sigma power (k= 9). Five (56%)[13,22,62–64] of nine[65–68] studies reporting on sex-based differences in sigma power observed higher absolute or relative sigma power in females on central channels, including one reporting this true for only fast sigma on central electrodes (14-16Hz[13]), and one for only slow sigma (11.25-13Hz[64]) on central electrodes, and one finding this only when considering relative slow sigma power (11.25-13Hz on C4[64]). One of the studies that did not find sex-based differences indicated that both males and females showed similar age-related reductions in sigma power[68]. Concerning spindle morphology, one article on spindle peak frequency (Hz) using power spectral analysis found females had higher peak frequencies, one confirming this for both fast and slow spindles[59].

##### Normal Sleepers: Slow Wave Activity

Twenty-four research papers explored sex-based differences in SOs (<1.25 Hz) and SWA (0.25-4.5 Hz) characteristics in NS. The 24 studies reported a total of 3,996 participants (mean age= 42.30 ± 18.30, median group size= 31, range 16 to 1595; see **Table 2**) and eight differing SWA characteristic outcomes either based on power spectral analyses in the delta (∼0.25-4Hz) or SO (<1.25Hz) frequency range or based on event detection (SW density, SW amplitude, SW frequency, SW slope, SW incidence, SW positive phase duration and SO event density).

##### Slow Wave and Slow Oscillation (event-based detection)

The most commonly reported domain was SW amplitude (k= 4). SW detection methods varied between papers (see **Table 2**). For SW amplitude, 75% (three of four studies[69–71]) found higher SW amplitude in females, while one did not[72]. Furthermore, one study found an age-by-sex interaction, showing that SW amplitude was significantly higher in older adults than in younger ones, with a steeper decline observed in older men[71]. Two studies reported steeper SW slope in females, with one finding this relationship regardless of age[71], and one finding females had a steeper SW slope across the lifespan[69]. SW frequency and SW positive and negative phase duration appear higher in males[70] (k= 1). Of three studies on SW density, one paper did not find sex-based differences[72], one found greater SW density in females[70], while the other found that with adapted criteria, older adults showed lower SW density than younger participants, regardless of sex[71]. There was no standalone sex effect, but a sex-by-age interaction emerged—older females retained stronger SW density, compared to older men. This suggests that while aging reduces SW features overall, females tend to maintain greater SW density later in life. The sole study on SO density (<1.25Hz) found that females exhibited higher SO density than males[22].

##### Delta and Slow Oscillation Spectral Power

The most frequently examined domain was delta spectral power (k = 21), followed by SO power (k= 3). Among the 21 studies on delta power, 9% (k= 2) reported no sex-based differences in relative delta power[64,73], and 48% (k= 10) reported no differences in absolute delta power[65,66,69,72,74–79]. Conversely, nine (43%) studies found higher absolute delta power in females[13,62,63,73,80–84], one (5%) in relative delta power[57], and one (5%) study[68] reported greater age-related reductions in absolute delta power in males within the 0.75–4.5 Hz range, but not when looking at mean delta power density in the broader 0.5–4.5 Hz range. Notably, one study[64] reported that this sex-based difference was limited to the SO range (0.75–1.25 Hz) and not observed in the 1.25–4.5 Hz range, while another[83] found it only during the first three NREM cycles. In contrast, one paper[73] found that when delta power was normalized to the percentage of delta activity in REM, females exhibited lower delta activity than males. Of the 10 studies (40%) that did observe sex-based differences in SWA, most focused on absolute delta power, rather than relative or normalized measures. One study[81] noted that elevated absolute delta power in females was most pronounced in the Pz and O2 channels. One study on SW incidence reported no differences between sex[72]. Finally, all three studies examining absolute and relative SO power (<1.25 Hz)[13,57,64] reported higher SO power in females compared to males (100%).

##### Summary

- Most studies (55%) investigating brain oscillations in NS did not examine sex-based differences.
- There is some evidence that females exhibited higher spindle density (25% of papers), sigma power (67% of papers), and spindle amplitude (67% of papers) compared to male NS. Measures of relative sigma power showed fewer differences.
- There were inconsistent findings on the presence of sex-based differences in SWA. SW amplitude appeared to be higher in females (75% of papers). Females also tended to exhibit higher absolute delta power than males (48% of papers), although half of the studies showed no difference in absolute delta power. Measures of relative delta power showed fewer differences.

#### Individuals with Insomnia

Seventeen studies on persons with insomnia only were eligible for the systematic review (See **Table 3**). Of those 17 papers, two directly compared sex-based differences in brain oscillations during sleep. All studies reported on primary insomnia using DSM-IV criteria (i.e., difficulties falling and/or staying asleep for at least one month).

##### Individuals with Insomnia: Spindle Activity

One paper explored sex-based differences in spindle characteristics in those with insomnia. They reported a total of 45 participants (mean age= 45.1 ± 7.5; **Table 3**) and one spindle characteristic outcome (i.e., sigma power).

*Spindle (event-based detection).* No studies to date have examined sex-based differences in spindle characteristics among persons with insomnia using event detection.

##### Sigma Spectral Power

A single study was conducted on sex-based difference in sigma power in IN[97]. Bader et al. (2013) delved into the investigation of potential sex-based differences in sigma power (12-16 Hz) during NREM 2 and 3 phases among individuals with insomnia. Following an initial adaptation night, the researchers reported no significant disparities between sexes, though specific statistical data were not provided, within a participant pool comprising mostly females (76%).

##### Individuals with Insomnia: Slow Wave Activity

Two research papers explored potential sex disparities in SWA characteristics in individuals experiencing insomnia, representing a total of 59 participants (mean age= 47.0 ± 8.7, median group size= 29.5, range 14 to 45; **Table 3**) and one SWA characteristic outcome (delta power).

##### Slow Wave and Slow Oscillation (event-based detection)

No studies to date have examined sex-based differences in SW or SO characteristics among individuals with insomnia using event detection.

##### Delta and Slow Oscillation Spectral Power

The sole reported domain was delta power (k= 2). Two papers investigated sex-based differences on delta spectral power, but found no significant results regarding absolute[97,98], nor relative delta power[97]. However, neither study reported statistical data. Both papers adhered to the DSM-IV criteria for insomnia disorder, conducted the research following an adaptation night, and tested absolute delta power. However, Bader et al. and Hall et al. employed distinct definitions for delta power frequency (0.1-3.5 Hz and 0.5-4 Hz respectively). No studies to date have examined sex-based differences in SO power among persons with insomnia using power spectral analysis.

##### Summary

- There were very few studies on brain oscillations in insomnia disorder. Most studies (88%) investigating brain oscillations in IN did not examine sex-based differences.
- There is some evidence that sigma and delta power in insomnia are not significantly affected by sex.
- No studies examined sex-based differences in spindle, SW and SO characteristics, or SO power in insomnia.

#### Insomnia and Normal Sleepers

Forty-four papers with both normal sleepers and persons with insomnia were included in the systematic review. Of those eight directly compared sex-based differences in brain oscillations during sleep (see **Table 4**). Out of the eight studies evaluating sex-based difference: eight reported on spindle characteristics, six reported on SWA characteristics, and one reported specifically on SO characteristics. The results are shown based on the main differences between the groups (NS vs. IN) and whether these differences depend on sex (i.e., sex by group interaction). When relevant data was provided, the overall effect of sex was also included. Half of the studies (50%) used DSM-IV criteria for primary insomnia, which define it as difficulty falling or staying asleep for at least one month without another underlying cause. A quarter (25%) used chronic insomnia criteria from DSM-III/IV/V, which require symptoms to persist for at least three months. A smaller number of studies (12.5%) focused on sleep maintenance insomnia (characterized by frequent nighttime awakenings; DSM-III) or paradoxical insomnia, where individuals report poor sleep despite objective evidence of adequate sleep (ICSD-3). Because so few studies used alternative diagnostic definitions, it’s difficult to determine whether the type of classification meaningfully influenced the findings.

**Table 4.**
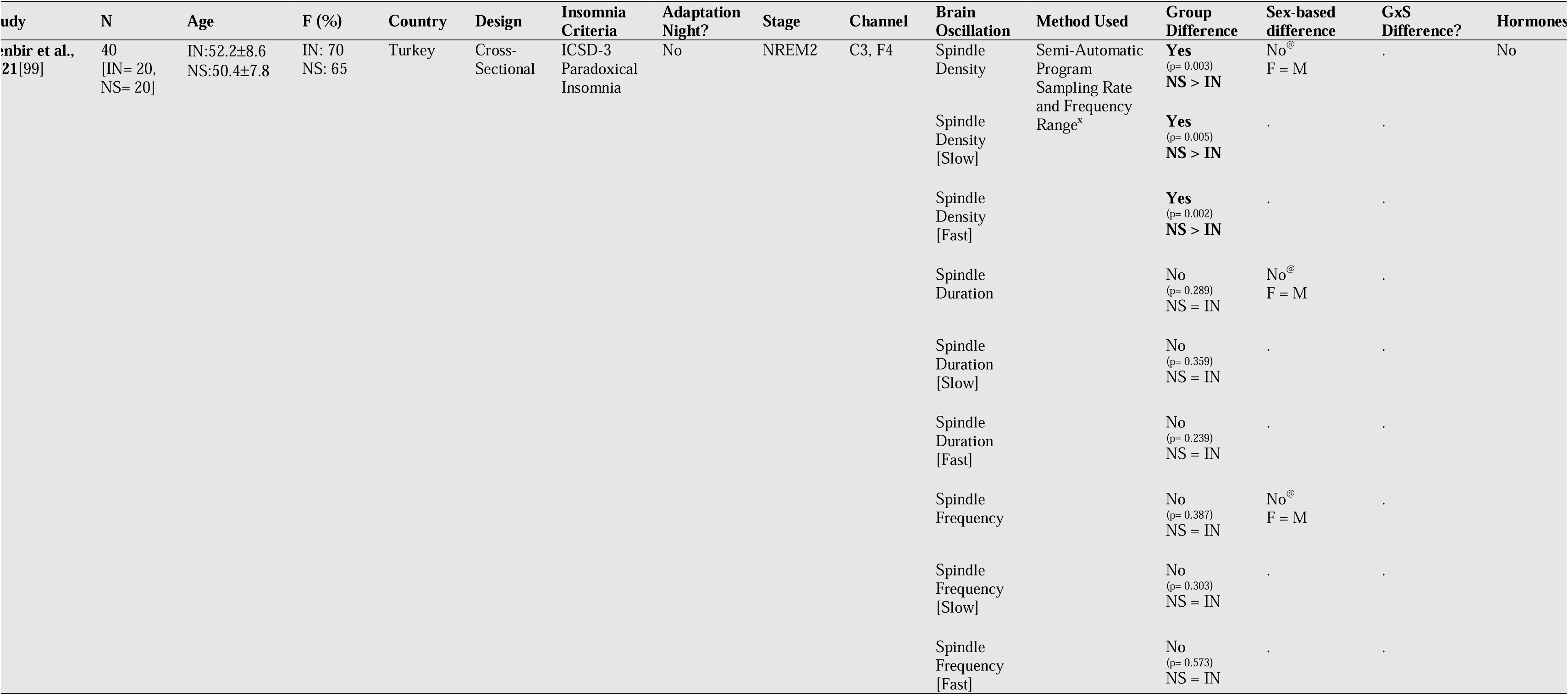
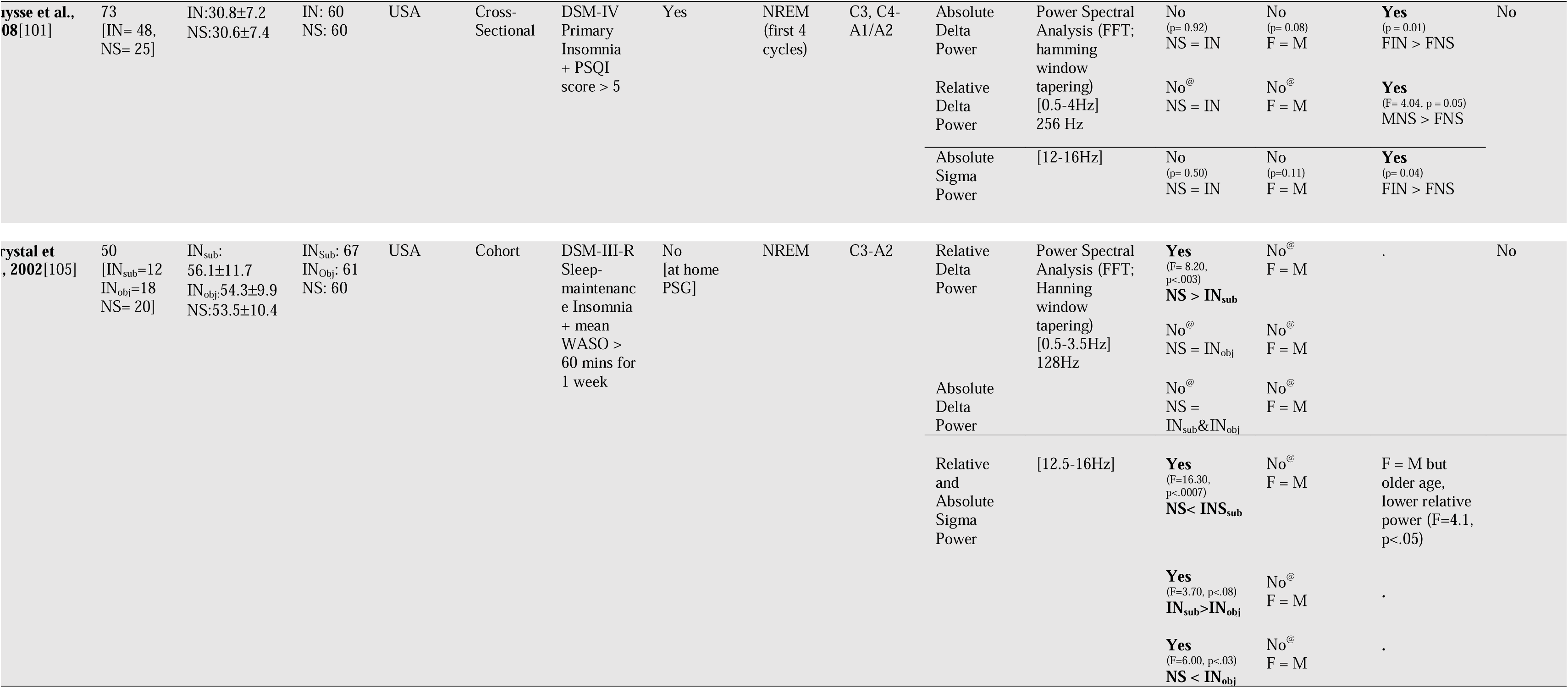
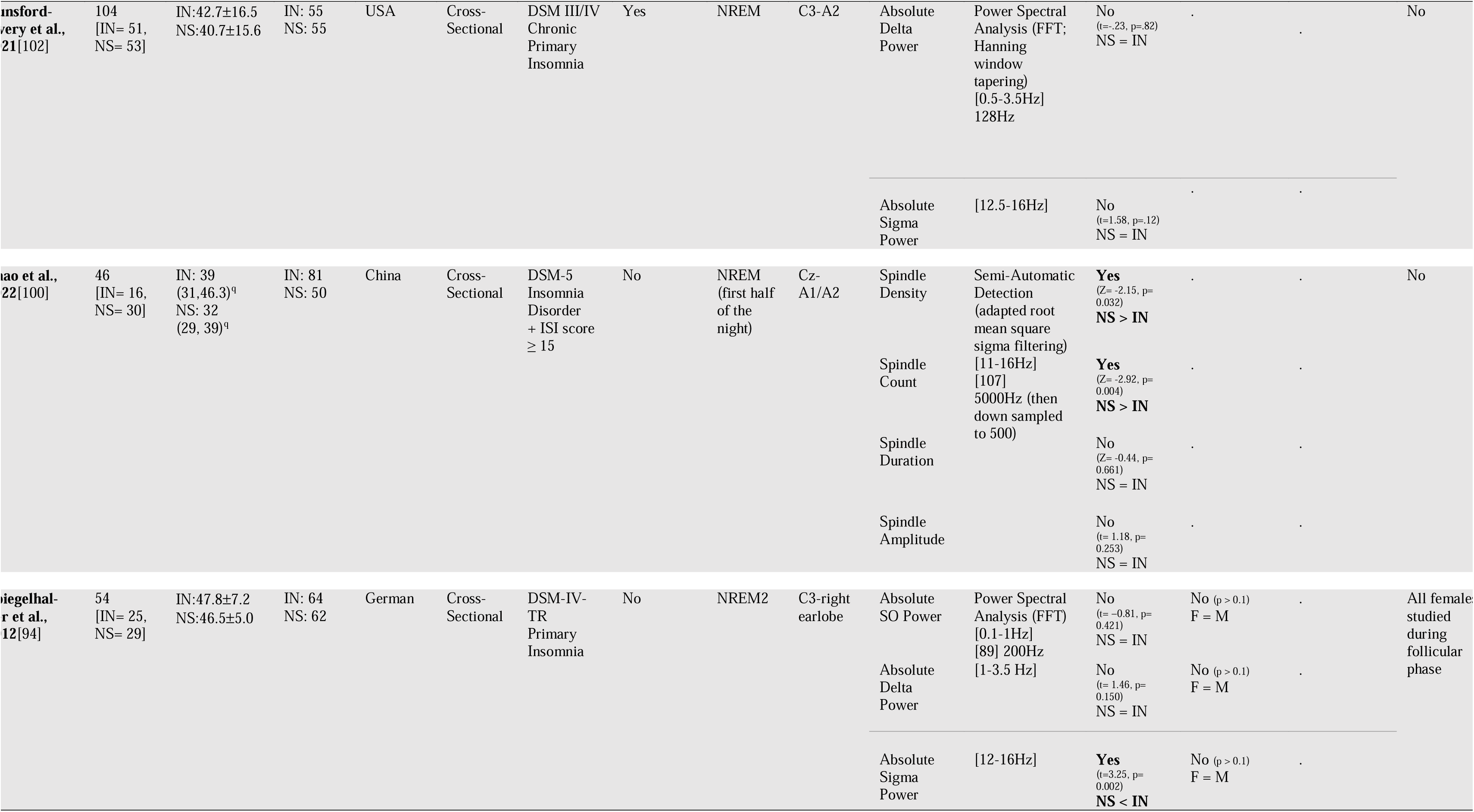
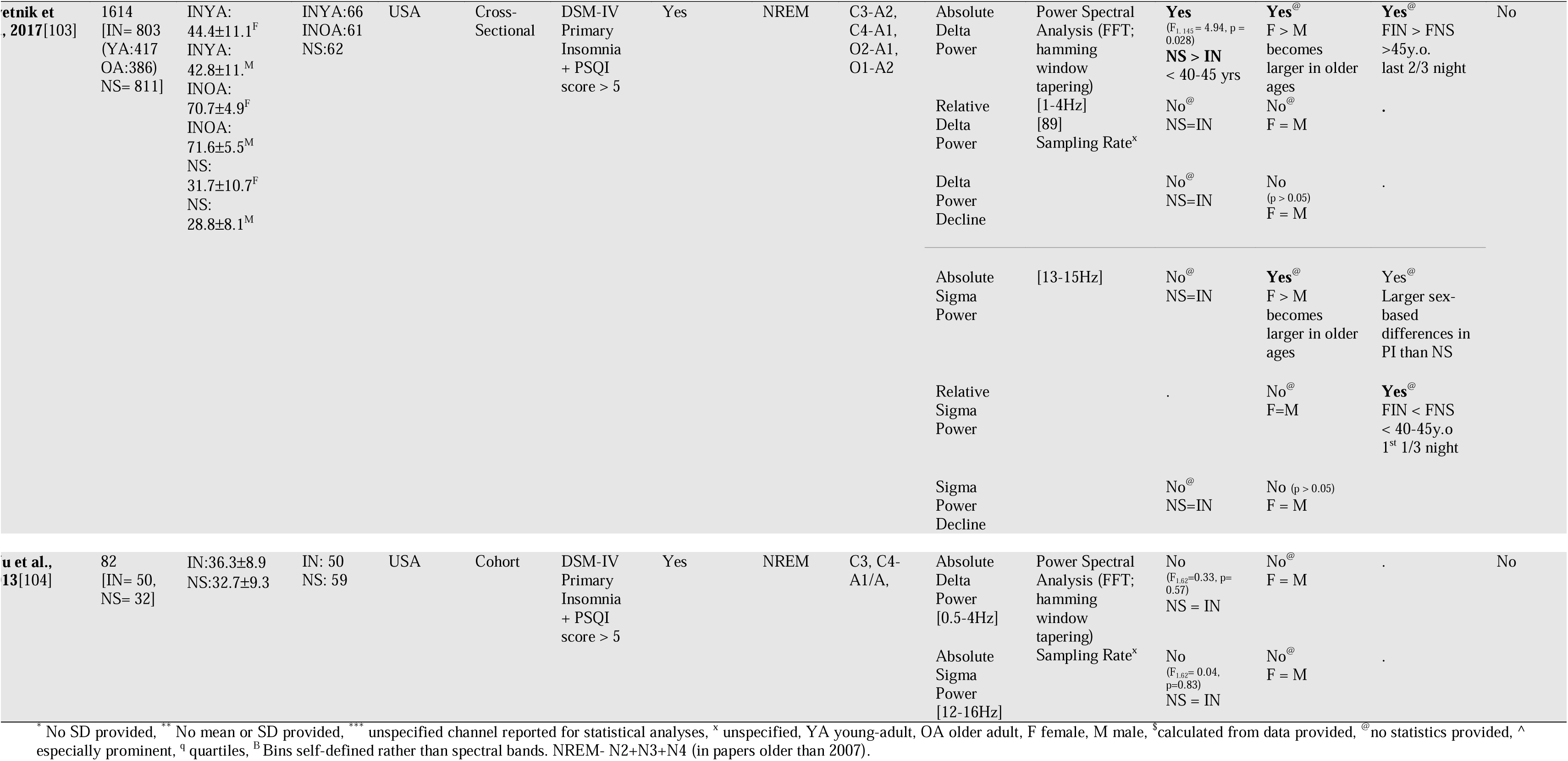
Study Characteristics of Insomnia and Normal Sleepers

##### Insomnia and Normal Sleepers: Spindle Activity

Eight research papers explored sex-based differences in spindle characteristics in normal sleepers and persons with insomnia, representing a total of 2,063 participants: 1,043 with insomnia (IN; mean age= 53.50 ± 9.20, median group size= 39, range 12 to 417) and 1,020 normal sleepers (NS; mean age= 40.90 ± 9.20, median group size= 29.50, range 20 to 811; See **Table 4**). There were six differing spindle characteristic outcomes (spindle density, frequency, amplitude, duration, count, and sigma power).

##### Spindle (event-based detection)

***Group:*** When investigating group effects without considering sex-based differences, the most reported domains were spindle density and duration (k = 2). Spindle detection method varied between papers (see **Table 4**). Neither studies reported any difference in spindle duration between NS and IN[99,100], even when further dividing spindles into slow or fast spindle activity (using Cz in the 11–16 Hz range and an unspecified range in C3/F4, respectively). The same studies found that NS had significantly higher spindle count[100] and density (combined, fast, and slow[99,100]) compared to IN. No group differences were found in spindle frequency[99] and amplitude[100]. ***Sex x Group:*** No studies have considered a sex*group interaction in any of the spindle characteristics. When considering the entire sample rather than by group, one study (50%; [99]) reported no significant sex-based differences on spindle density, duration, and frequency, but did not report statistical data. The other paper on spindle duration, count and amplitude only compared groups (IN vs. NS) without analyzing sex-based differences[100].

##### Sigma Spectral Power

***Group:*** When investigating group effects without considering sex-based differences, sigma power was the most commonly reported domain in power spectral analysis (k = 7). Of the studies examining group differences in sigma power in NS and IN, 57% (k= 4)[101–104] found no significant differences between groups for absolute and relative power. Two studies (29%)[94,105] found that IN had greater absolute and relative sigma power compared to NS, with one of the studies finding this to be true for both subjective and objective insomnia[105]. Specifically, those with subjective insomnia (previously called paradoxical insomnia-[106]) had significantly higher sigma power than objective insomnia (previously categorized as psychophysiological insomnia) and NS[105]. The sole study on sigma power decline across the lifespan[103] reported no group differences in all parts of the night. This indicates that the rate of absolute sigma power decline with age was similar for both IN and NS groups, and throughout all parts of the night. ***Sex x Group:*** When investigating sex effects, six studies examined sex-based differences in sigma power in NS and IN. Of those, 83% (k= 5)[94,101,103–105] found no significant effects in either absolute or relative power, with only 20% providing statistical support[94,101]. Moreover, two studies reported a group-by-sex interaction.

One found that female IN had significantly greater absolute sigma power than female NS, while no significant differences were observed between male IN and male NS, nor were there sex-based differences within either group[101]. The other study reported that although there were no group differences in absolute sigma power, there were larger sex-based differences within the IN group than NS, and the sex effect was more apparent in older ages[103]. Conversely, they found that female NS had significantly higher relative sigma power than female IN, specifically in those under 40–45 years old during the first third of the night[103]. The remaining article only compared groups without analyzing sex-based differences in sigma power[102]. The sole study on sigma power decline across the lifespan[103] reported no group or sex-based differences in all parts of the night, but did not assess group-by-sex effects. This indicates that the rate of absolute sigma power decline with age was similar for both females and males, across IN and NS groups, and throughout all parts of the night.

##### Insomnia and Normal Sleepers: Slow Wave Activity

Six research papers explored sex-based differences in SWA characteristics in normal sleepers and individuals with insomnia. The six studies reported a total of 1,977 participants: 1,007 with insomnia (IN; mean age= 42.50 ± 10.10, median group size= 49, range 25 to 803) and 970 normal sleepers (NS; mean age= 40.80 ± 9.50, median group size= 30.5, range 20 to 811; See **Table 4**) and two differing SWA characteristic outcomes (delta and SO power).

##### Slow Wave and Slow Oscillation (event-based detection)

No studies to date have examined group or group-by-sex-based differences in SW or SO characteristics using event detection.

##### Delta and Slow Oscillation Spectral Power

***Group:*** When investigating group effects without considering sex-based differences, the most commonly reported domain was delta power (k= 6). Of papers on delta spectral power, 67% (k= 4) found no group differences in relative[101] or absolute[94,101,102,104] delta power, while 29% (k= 2) found a group difference—one for relative but not absolute[105], and the other for absolute but not relative[103]. One study (14%) found that IN had higher absolute SWA compared to NS, specifically in those under 40–45 years old during the first third of the night[103]. The other (14%) found the opposite, where NS had greater relative SWA compared to subjective IN, but not objective IN[105]. Similarly, one paper investigating SW power decline (i.e., across the lifespan) reported no group differences in all parts of the night[103]. This indicates that the rate of absolute power decline with age in the delta band was similar across IN and NS groups, and throughout all parts of the night. The only study on SO power (<1.25 Hz) found no statistically significant differences between groups[94]. ***Sex x Group:*** When investigating sex effects, the most commonly reported domain was delta power (k= 6). Of papers on delta spectral power (k= 6), 83% (k= 5) found no sex-based differences in relative[101,103] or absolute[94,101,104,105] delta power, while one (14%) found sex-based differences[103] that became more apparent in older ages. Specifically, they found females had greater absolute SWA, but no differences in relative delta power. Two studies (29%) investigated group-by-sex interaction and both found that female IN had higher absolute delta power compared to female NS[101,103]. However, Sventik and colleagues (2017) found this relationship specifically in females older than 45 years and in the last two-thirds of the night, whereas Buysse and colleagues (2008) found this for the full NREM period in their middle-aged female population. A group-by-sex interaction was not observed on relative delta power in both of these studies. However, one found[101] that male NS had higher relative delta power compared to females (a contrast to the findings in our NS only section). Similarly, one paper investigated delta power decline (i.e., across the lifespan) reported no sex-based differences in all parts of the night[103], but did not assess group-by-sex effects. This indicates that the rate of absolute power decline with age in the delta band was similar for both females and males, and throughout all parts of the night. The only study on SO power (<1.25Hz) found no statistically significant differences between sexes and did not examine group-by-sex interactions[94].

##### Summary

- Most studies (80%) investigating brain oscillations in IN and NS did not examine sex-based differences (i.e., perform a main effect of sex or group-by-sex interaction analyses).
- No clear or consistent sex-based differences were found in spindle characteristics although some studies showed slight decreases in sigma power, particularly in older females with insomnia, but results were inconsistent.
- Studies investigating SWA also showed mixed results, with some studies finding higher delta power in females with insomnia, particularly in middle (23-38 years) and older (> 45 years) age groups.
- Overall, findings on sex-based differences in sleep-related brain oscillations in individuals with and without insomnia were inconclusive and varied by study method, age, and analysis type.

### Findings From the Meta-Analysis

While there were enough eligible papers and statistical data for meta-analysis of sex-based differences on brain oscillations in normal sleepers, there was insufficient statistical data provided by papers to meta-analyse the effect of sex on brain oscillations in insomnia compared to normal sleepers.

#### Overall Effect of Sex on Spindles in Normal Sleepers

Two[58,59] of the eight studies on spindle density in normal sleepers were eligible for meta-analysis, revealing a moderate sex-based difference effect, favouring females (n= 190, SMD -0.65 [95% CI: -1.95, 0.66], I^2^= 85% - **Figure 2A**). Of the eight studies on absolute and normalized sigma power in NS, five provided enough data to allow meta-analysis of absolute sigma power[13,66–68,101], resulting in a statistically significant but small effect size (n= 365, SMD -0.29 [95% CI: -0.50, -0.08], I^2^= 0% - **Figure 2B**) favouring females. There were not enough papers reporting data for meta-analysis on the effects of sex on other spindle characteristics (e.g., relative sigma power, spindle density and duration).

**Figure 2.**
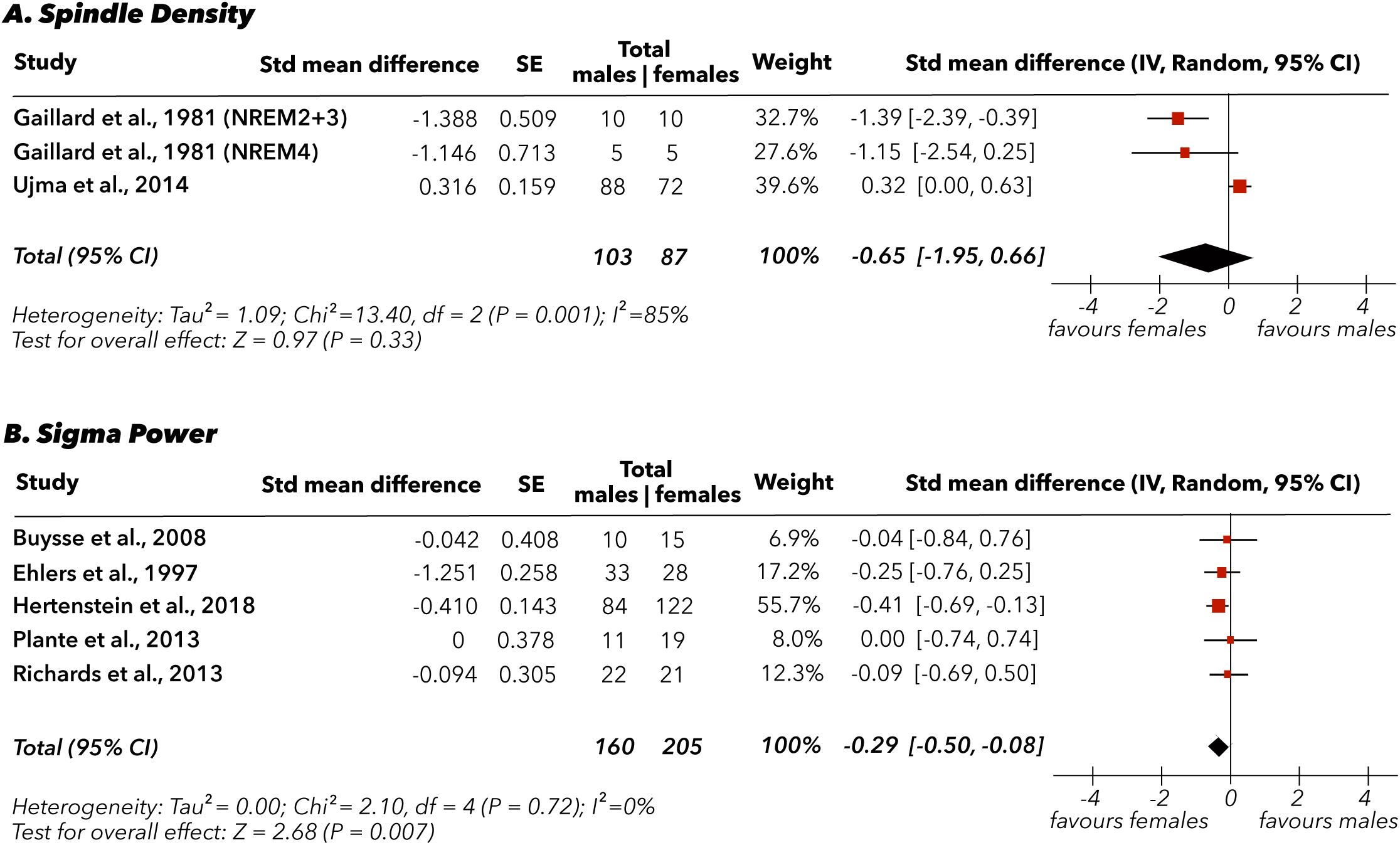
Synthesis and forest plot for spindle outcomes [(A) spindle density; (B) sigma power)] of male vs female. CI, confidence interval; SE, standard error; Std., standardised.

#### Overall Effect of Sex on Slow Wave Activity in Normal Sleepers

Seven of the 19 studies on delta power were eligible for meta-analysis. The analysis of six studies (**Figure 3**) on absolute and one on relative delta power in normal sleepers showed a statistically significant difference, favouring females with a moderate effect size (n= 407, SMD - 0.50 [95% CI: -0.70, -0.30], I^2^= 0%). There was insufficient data provided by papers to meta-analyse the effect of sex on SW and SO (based on event detection) characteristics.

**Figure 3.**
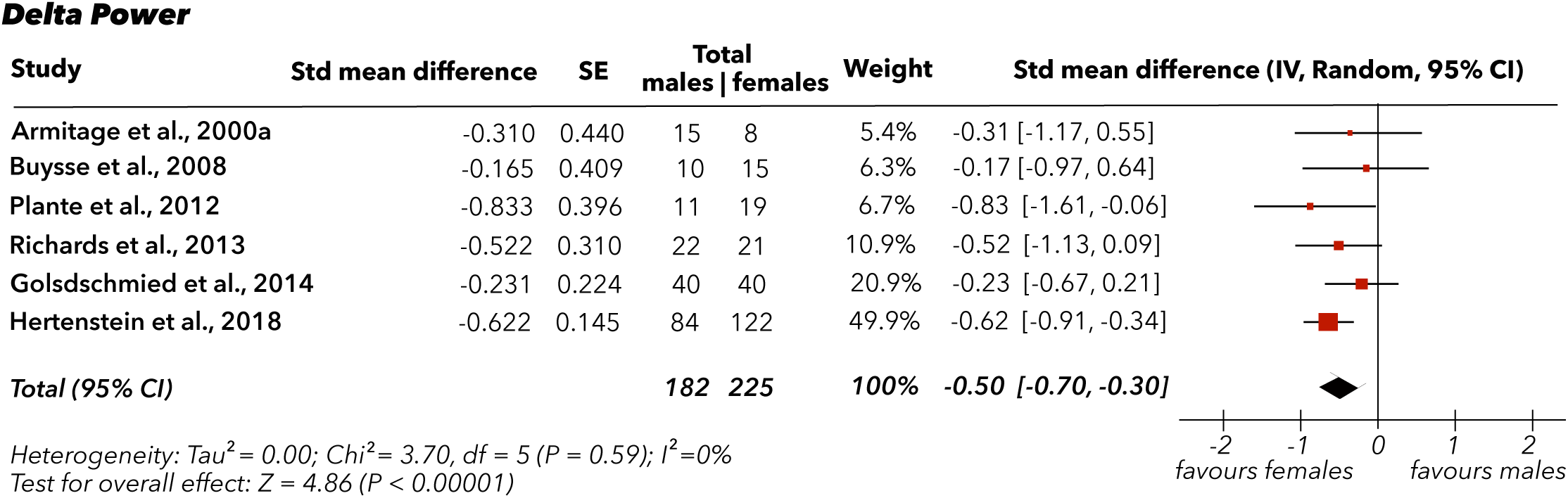
Synthesis and forest plot for delta power outcomes of male vs female. CI, confidence interval; SE, standard error; Std., standardised.

### Heterogeneity

Heterogeneity across studies was observed, particularly in the reporting of spindle and SWA characteristics. Significant heterogeneity was found across studies for sleep spindle density (I²= 85%), but there was no observed heterogeneity among sigma power (I²= 0%) or delta power (I²= 0%).

We conducted several **sensitivity analyses** to further investigate heterogeneity and the robustness of our results. We performed **leave-one-out analyses**, systematically excluding one study at a time to evaluate if any single study unduly influenced the results. None of the individual studies appeared to significantly alter the direction or magnitude of the effects. Finally, we conducted sensitivity analyses to assess the impact of **small-study effects** using funnel plots, which showed no significant asymmetry, suggesting no major concerns regarding publication bias or small-study bias in the results.

We assessed **publication bias** visually using funnel plots of effect size (Hedges’ g) by standard error to identify any potential reporting bias in the studies. No significant asymmetry was observed in the funnel plots, suggesting that there was no apparent publication bias for the outcomes of interest (i.e., sleep spindle density, sigma power, SWA).

## DISCUSSION

To the best of our knowledge, this is the first systematic review and meta-analysis to examine the effects of biological sex on sleep spindles and SWA in adults with and without insomnia and to directly assess whether insomnia modifies these sex-related patterns in quantitative EEG measures (e.g., event-based detection, power spectral analysis). We found small to moderate effects for higher spindle density, sigma power, and delta power among females compared to male normal sleepers. Our review also highlights critical gaps in the literature for sex-based differences in brain oscillations during sleep.

### Sex-Based Differences in Spindle and Slow Wave Activity in Normal Sleepers

Sleep spindles and SWA are key oscillatory features of NREM sleep, involved in synaptic plasticity, memory consolidation, and cognitive regulation[24,25,31,32]. Evidence from the systematic review and our meta-analysis found modest but consistent sex-based differences in both spindle and SWA characteristics, generally favouring females. Few studies found sex-based differences in ***spindle*** density[58,59], but those that did typically reported higher values in females[57,59]. Our results confirmed a moderate female advantage in spindle density, most rated good to satisfactory quality. All five papers that failed to detect sex differences had small sample sizes, likely a result of insufficient statistical power, analyzed spindles only during NREM2[45,46,55,56,108], and had inconsistent methods and protocols (e.g., EEG channels, sigma frequency definitions, sampling rates, participant age, use of an adaptation night; see **Table 2**). Although the evidence for other spindle characteristics (e.g., amplitude, duration, and distribution) was more variable and not suitable for quantitative analyses, the overall pattern suggests a modest female advantage in several spindle-related features. This aligns with findings from the influential study by Purcell et al. (2017)[109], which was excluded from this review due to inclusion of children and persons with sleep apnea in a pooled dataset.

Most studies reported higher absolute and relative sigma power in females[13,57,62–65], particularly in the during NREM and in central channels Our results confirmed a small but consistent female advantage in absolute sigma power. Those that failed to find significant sex-based differences in sigma power[43,65,66,68] may be due to insufficient statistical power due to limited sample sizes. While relative sigma power could not be included in the meta-analysis due to an insufficient number of studies, Chapman et al. (2025)[110] reported moderate certainty for no sex-based differences in middle to older adults. This contrasts with findings from our systematic review indicating that females display greater relative spindle power compared to males[57,64]. This may be due to their inclusion of papers using non-gold-standard methods (e.g., at home PSG).

For ***SWA***, most studies examined delta power (0.5–4 Hz), with fewer focusing specifically on SW (<4 Hz) or SO (<1.25 Hz) as discrete events. Our systematic review and meta-analysis found consistent sex differences in delta power, with females generally showing higher absolute values[13,57,62,63,73,77,78,80–84]. Some studies highlighted regional effects in parietal and occipital areas [74]. These studies were mostly of good or satisfactory quality, and the results across studies were very consistent. Studies reporting no sex differences[64–67,75,77,79] in absolute delta power often had small samples, older participants, and lacked an adaptation night. In contrast, studies of relative delta power reported no sex-based differences[73,76,78–80]. This divergence between absolute and relative power likely reflects methodological and anatomical influences rather than genuine neurophysiological sex differences in delta power. Absolute EEG power is strongly affected by factors such as skull thickness and head tissue conductivity, which impact EEG amplitude and differ on average between males and females [63,71,110, 111]. By scaling power values within each individual, relative metrics reduce the influence of skull and tissue differences and improve comparability across individuals. The absence of sex effects in relative delta power therefore suggests that the differences observed in absolute power may be driven more by global amplitude differences (i.e., all power bands) than by frequency-specific neural processes. This has been supported previously by Chapman et al.’s (2025) review[112]. One study found greater normalized delta power in males[73] likely due to atypical normalization procedures. Support for sex-based differences in event-based findings on SW and SO features were too inconsistent and heterogeneous for quantitative synthesis, though some studies reported greater SW amplitude in females[69–71].

Our findings on SWA and spindle characteristics suggest potential sex-related differences in thalamocortical connectivity, which plays a crucial role in regulating cognitive functions[32], emotions and has implications on neuroprotection[113]. Across studies, we observed a stronger convergence in findings on spectral power (i.e., sigma and delta power) compared to event characteristics (i.e., SW and spindle events). This may reflect methodological consistency of spectral analysis which is more sensitive to sampling variability and detection algorithms.

### The Effects of Insomnia on Sex-Based Differences in Spindle and Slow Wave Activity

There was notable heterogeneity in how insomnia was defined across studies, which may influence the interpretation of results. The findings presented mainly represent the DSM-IV criteria for primary insomnia (requiring at least one month of sleep difficulty [50%]), others employed more stringent definitions, such as DSM-III/IV/V criteria for chronic insomnia (25%), which require symptoms to persist for at least three months. A small subset (12.5%) focused on specific subtypes, such as sleep maintenance insomnia or paradoxical insomnia. Given the limited number of studies using these alternative classifications, it remains unclear whether differences in diagnostic criteria significantly impacted the findings.

**Group Differences.** We found inconsistent findings for whether ***spindle*** activity remains stable regardless of insomnia. A small number of studies reported spindle density is more sensitive to the presence of IN[99,100], supporting existing evidence that reduced spindle density is a reliable marker of disrupted sleep[32,114,115]. Contrastingly, two studies found higher sigma power in IN compared to NS[94,105], while most found no differences in sigma power, spindle duration, frequency, and amplitude. While spectral measures like sigma and delta power offer stable, trait-like insights into sleep physiology in normal sleep, they may also miss subtle disruptions seen in insomnia and could explain findings of altered spindle density in insomnia despite inconsistent spectral findings. In contrast, event-based metrics such as spindle density, may better detect state-dependent disturbances in thalamocortical and cortical function[31]. However, the inconsistent study quality and risk of bias may have influenced these findings[94,105].

To the best of our knowledge, no studies investigating sex-based differences have also examined differences between insomnia and normal sleepers in event-based SW or SO characteristics. Our systematic review also found limited but generally consistent evidence regarding ***SWA*** in insomnia. Specifically, the majority of studies found no significant differences between IN and NS in either absolute[94,101,104] or relative[101,102] delta or SO power (<1.25Hz)[94]. Our findings contrast with other studies[116,117] and systematic reviews/meta-analysis[118] that report lower absolute and relative delta and SO power in persons with insomnia compared to normal sleepers. This may be due to differences in methods and poorer quality of the studies (e.g., limited generalizability, potential bias from small or unrepresentative samples and non-responders, inadequate control for confounders, and questionable statistical validity) investigated at both group and sex differences. This might also underscore the heterogeneous nature of insomnia[119]. Subjective complaints may not align with objective measures (reflecting sleep state misperception[106,120]) while others may show altered sleep homeostasis (e.g., short sleep duration subtype[121]). However, most studies lack detailed reporting on insomnia duration (i.e., chronicity of symptoms) or subtype, which might have more profound impacts on neurophysiology. Emerging evidence also suggests that EEG profiles vary across subtypes. For instance, one study found that persons with subjective IN exhibited significantly higher sigma power compared to those with objective IN and NS[105].

**Sex-Based Differences.** Six studies tested sex-based differences in absolute and relative ***sigma*** power in IN[94,101–105]. Both studies on relative sigma power found no sex differences. However, five reported no sex-based differences in absolute sigma power[94,101,102,104,105], one reported higher absolute sigma power in females when combining IN and NS groups[103], while the other found no sex-based difference in the IN group alone[97]. Only two studies tested for sex-by-group interactions. One[101] found that middle-aged female IN exhibited greater absolute sigma power than NS, while the other reported greater absolute sigma power in females than males, with sex differences more pronounced in the IN group than in NS[103]. The opposite pattern was found for relative sigma power[103]—lower in female IN, specifically during the first third of the night. These discrepancies likely arise from differences in methods—such as using absolute versus relative power, electrode placement (central vs. lateral), insomnia subtypes, and time-of-night studied. Studies that found no sex effects in sigma power[94,104,105] or other spindle characteristics [99,100], often combined insomnia and normal sleeper groups or did not test sex or group-by-sex interactions systematically. Compared to those that did find significant sex effects, some of these had lower quality, limiting the reliability of their findings[94,105].

Research on sex-based differences in ***SWA*** and SO features in insomnia is even more sparse and inconclusive, with no studies, to the best of our knowledge, systematically examining sex differences or sex-by-group interactions using event detection methods. Of the few that examined SWA in insomnia, only one study identified sex-based differences in absolute delta power[103], with a female advantage that became more pronounced with age. Every other study reviewed found no significant sex effects in either relative[97,101,103] or absolute[94,97,98,101,104,105] delta power. Compared to the one study finding significance, most of these studies were of unsatisfactory quality, limiting the reliability of their findings and making it difficult to draw firm conclusions[94,104,105]. Only two studies investigated group-by-sex interactions in delta power; both found that female IN had higher absolute delta power compared to female NS[101,103], one specifically noting this in older female IN (>45 years). However, one[101] reported higher relative delta power in male NS compared to females—a finding that contrasts with other studies showing no sex differences in normal sleepers. This discrepancy may reflect population differences (e.g., age, hormonal status), methodological factors, or an artificially inflation from using relative power. In this case, higher relative delta power in males may not reflect stronger SWA, but rather lower overall EEG power compared to females, making delta proportionally larger.

Given the quality concerns (i.e., high risk of bias, small samples, and methodological inconsistencies), and heterogeneity in insomnia criteria, it remains difficult to conclude whether sex-based differences are present in NREM oscillatory activity in insomnia disorder. Systematic reviews reinforce these findings, noting that while sex-based differences in spindle and SWA are robust in normal sleeping populations (i.e., mostly favouring females), they are not consistently replicated in clinical insomnia samples[37].

### Unraveling the Female Insomnia Paradox

Females generally exhibit higher spindle density, SWA, and longer total sleep time—features typically associated with deeper, more restorative sleep. Yet, they are nearly twice as likely as males to develop insomnia disorder[1]. Although healthy females typically show favorable sleep features, it is unclear whether these are preserved in females with insomnia, as few studies have examined group-by-sex interactions and female-specific group analyses. These physiological advantages may be diminished in insomnia or may not fully protect against stressors and stress reactivity more prevalent in females—such as hormonal fluctuations (e.g., menopause), caregiving demands and may further heighten vulnerability to sleep disruption[122].

Additionally, standard EEG measures may not capture subjective sleep experiences or arousal regulation, both of which contribute to insomnia. Thus, this paradox underscores the need to examine sex-specific sleep physiology across normal and clinical sleep, integrating biological rhythms with psychosocial context and lived experience.

### Layers of Influence: Age as the Thread Connecting Sex, Hormones, and Insomnia

#### Insomnia

Insomnia prevalence and manifestation are highly dependent on both age and sex given the role of hormonal changes across the lifespan. Indeed, sex disparities in sleep only appear at puberty, and the risk for developing insomnia increases nearly threefold after menarche, and continues to spike during key reproductive transitions, such as pregnancy, peri-and post-menopause[8,9,123]. During the menopausal transition and reduction in ovarian hormones, insomnia prevalence rises from 35% to 53%[9,124]. Conversely, menopausal hormone therapy (MHT; i.e., bioidentical or low-dose estrogens and progestogens) has been shown to improve sleep quality in perimenopausal females[125,126,127], while hormonal contraceptives (i.e., synthetic ethinyl estradiol and progestins) report more severe insomnia symptoms than non-users[128]. Changes in male gonadal steroid levels across the lifespan (e.g., andropause, anabolic androgenic steroids use), are also associated with poorer sleep quantity and quality[125,129,130]. These findings underscore age as a central thread- shaping hormonal transitions, modulating the brain’s sleep-related oscillatory activity, and showing a consistent, linear relationship with sleep complaints. In this way, age exerts a powerful and consistent influence shaping, amplifying, or masking the effects of sex-based differences on sleep.

**Age.** Age is a critical factor that underlies variability in sex-based differences in spindles and SWA across normal and insomnia populations. Both spindle characteristics and SWA decline significantly with age. It has been shown that ***spindle*** activity peaks in early adulthood and diminishes with aging due to structural and functional brain changes. This includes normal neuronal loss, reduced thalamocortical connectivity, diminished GABAergic neurotransmission, and cortical thinning[109,131,132]. Similarly, ***delta*** power declines gradually throughout adulthood, reflecting neurobiological deterioration-particularly in the frontal cortex-rather than altered sleep patterns alone. These age-related changes reduce the brain’s capacity for deep, restorative sleep, leading to shallower, fragmented sleep and contributing to the increased prevalence of insomnia in older adults[132,133]. Spindle and SWA are also linked to cognitive health, raising the stakes for understanding their modulation by age and insomnia. For instance, reduced spindle and SWA are associated with cognitive impairment and increased risk for Alzheimer’s disease (AD)[134,135]. Since females represent approximately two-thirds of AD cases, with estrogen decline exacerbating this trend, understanding how age and sex interact to shape sleep oscillations are key to both sleep health and cognitive long-term cognitive trajectories. Insomnia emerging in midlife is also linked to subjective cognitive decline, a potential early sign of mild cognitive impairment (and possible dementia), and associated with objective memory decline in males[136,137]. Few studies in this review directly examined age-by-sex interactions; however, some evidence suggests that differences in spindle and SWA between individuals with insomnia and normal sleepers may vary across the lifespan. For example, Svetnik et al. (2017)[103] found significant group-by-age effects in females only: younger and middle-aged females with insomnia (<40 years) showed lower relative ***sigma*** power compared to normal sleepers, while older females with insomnia (>45 years) showed greater absolute ***delta*** power than age-matched controls. These findings suggest that the impact of insomnia on sleep microarchitecture may evolve with age in females, potentially reflecting age-related neurobiological changes. Given the wide age range in our review (23.6–77.1 years) and the limited number of eligible studies, age-stratified analyses were not feasible, complicating interpretation of age-related effects.

#### Sex Hormones

Hormonal changes across the lifespan likely shape sex-specific patterns in sleep physiology. Sex-steroid hormones (i.e., estrogen, progesterone, and androgens), while primarily are important for maintaining hormonal homeostasis, also play complementary roles in supporting sleep by influencing brain regions involved in sleep regulation (e.g., modulating thalamocortical oscillatory activity). This can be observed in rodent studies which demonstrate that removing gonadal steroids eliminates sex-based differences in sleep architecture[138,139]. Estradiol and progesterone help quiet brain activity by enhancing GABAergic inhibition in areas that generate sleep ***spindles*** (i.e., thalamic reticular nuclei and cortical layers), making these rhythms more likely to occur[140,141]. For example, spindle density increases during the luteal phase (high estradiol and progesterone) and with oral contraceptive use[142], while combined hormone replacement therapy (HRT) or single-hormone regimens are associated with higher spindle activity in pre and postmenopausal females[140,143,144]. Testosterone may also modulate spindle activity, but the evidence is mixed and less consistent. Animal studies show that gonadal steroid injections in castrated males and ovariectomized females eliminate sex-based differences in sleep spindle expression implying an interaction between androgens and underlying sex-based neural circuitry[145]. In humans, testosterone fluctuations (such as in aging men or women with polycystic ovary syndrome) may alter spindle characteristics given the impacts on poor sleep quality, but this remains understudied[146].

While ***SWA*** appears less sensitive to hormonal fluctuations, likely due to its regulation by sleep homeostasis, some evidence suggests hormones still play a modulatory role. In animal models, estradiol suppresses delta power by acting on sleep–wake regulatory regions (e.g., basal forebrain and hypothalamus)[139,145,147], yet human studies show little change in SWA during peri-menopause despite steep declines in estradiol[148]. This may possibly be due to gradual hormonal changes and compensatory neuroadaptation that may preserve sleep homeostasis despite estradiol fluctuations [145,147]. It has been hypothesized that progesterone could enhance deep sleep, however progesterone’s effects on SWA are inconsistent. Some studies report increases in delta power when progesterone is naturally high and others show reductions or no change [149,150]. Androgens on the other hand may influence SWA more directly by regulating sleep homeostasis. Testosterone acts on androgen receptors in cortical and subcortical structures, with particular influence on the frontal cortex—a key region for SWA generation[151]. In animal studies, testosterone depletion (e.g., gonadectomy) reduces SWA, while testosterone replacement restores it[138]. In humans, however, higher testosterone levels in males either endogenous or via therapy (e.g., in males with congenital GnRH deficiency) are linked to reduced SW sleep and delta power in males, even after controlling for age and other covariates[145,152]. Findings in females remain inconsistent[153], and andropause and SWA have not been directly studied. Overall, estradiol and progesterone enhance spindles via GABAergic mechanisms, but their effects on SWA are mixed or neutral. Testosterone may influence cortical structures involved in SWA, but it’s effects on spindle and SWA remain understudied.

Hormonal fluctuations seem to directly and indirectly effect sleep quality. Yet, the majority of studies (34 studies reviewed in this paper [80%] do not consider key hormonal variables that can modulate EEG activity. Of the 42 studies reviewed, only eight explicitly controlled for hormonal factors affecting sleep or EEG[55,63,66,68,70,72,73,94]. These controls included excluding hormonal contraceptive or HRT users, selecting based on menopausal status or FSH levels, and timing data collection to specific menstrual phases (e.g., follicular phase). Some also ensured participants were not menstruating or reported contraceptive use.

### Lack of Sex-Based Research in Sleep Brain Oscillations

Despite growing recognition of sex as a key biological variable, our review demonstrates that sex-based differences in spindle and SWA characteristics are infrequently and inconsistently examined in sleep research. While some accounted for sex through design elements like matched controls or covariates, few examined it directly (See **Supplementary Information**). Even of the sex-based difference studies, sex was a secondary variable of interest. While most studies met basic methodological standards, only 5% were high quality, and nearly one-fifth were unsatisfactory. Common issues included small, sex-imbalanced samples, lumping data from both groups, lack of statistical support for sex effects, and failure to test sex-by-group interactions—which according to simulation studies, can reduce detection power of sex-specific effects by up to 28% and reduce reliability[154]

### Strengths, Limitations and Future Direction

Our systematic review and meta-analysis have several strengths. We used comprehensive search strategies without language or date limits, addressing a key limitation of prior reviews. By examining sex differences in sleep spindles and SWA in both normal sleepers and persons with insomnia, we offer a broader view of how biological sex influences brain rhythms during sleep (See **Figure 4**). Uniquely, we also explored how sex-steroid hormone fluctuations interact with thalamocortical synchronization to shape insomnia vulnerability. This mechanistic perspective not only clarifies sex differences in sleep disorders but also points to potential biomarkers for personalized treatments—such as targeting spindle deficits in stress-prone females or restoring SWA in perimenopausal women. By connecting neural activity, hormonal changes, and subjective sleep experience, our review advances a more biologically grounded understanding of sex-specific insomnia and highlights key gaps in the current evidence base.

**Figure 4.**
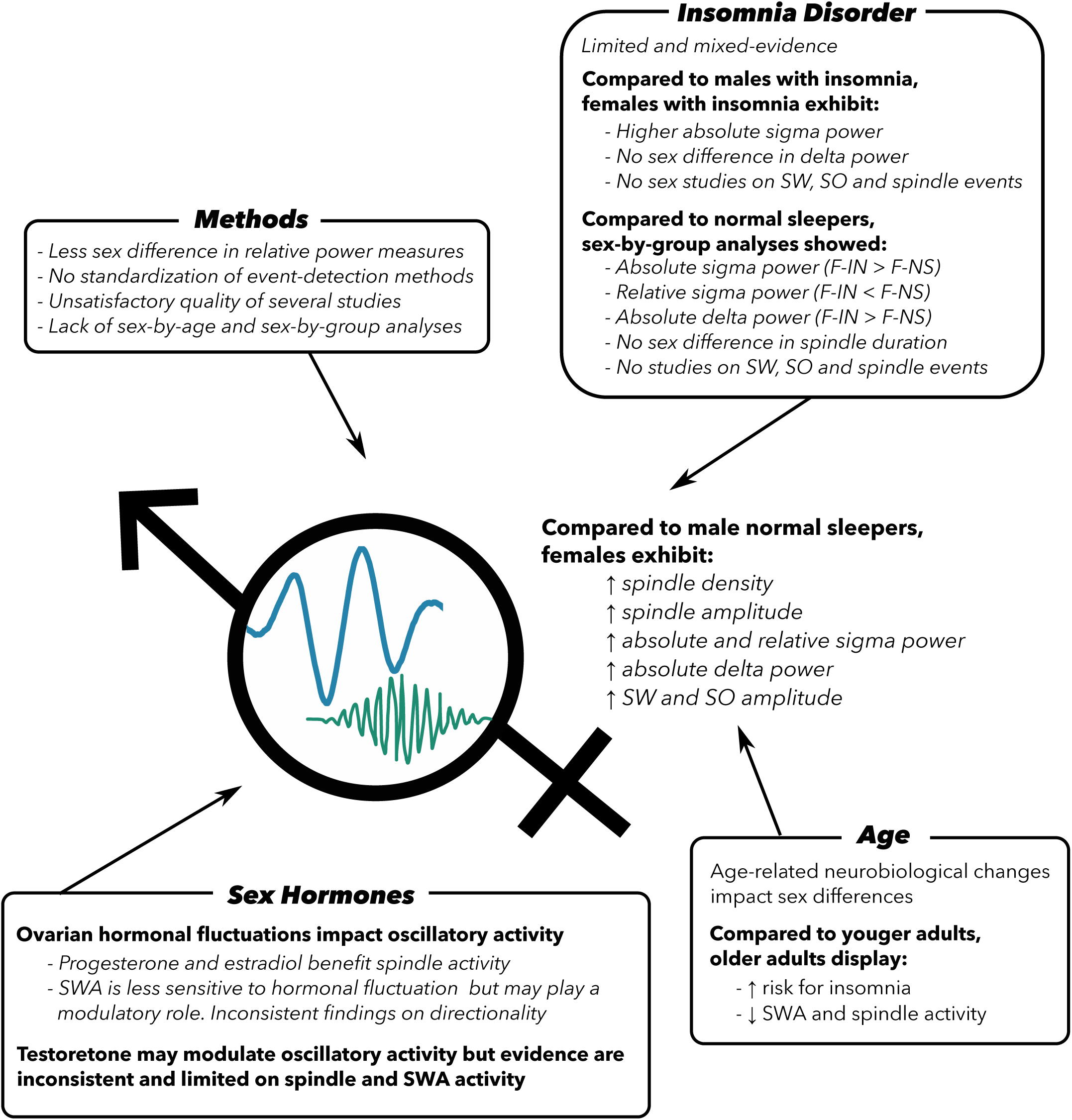
Overview of research on participant and study design factors affecting sleep oscillations

However, our systematic review also had limitations. The limited number of available studies, considerable variability in study methods, incomplete data (e.g., missing effect estimates or sample characteristics), and lack of information on the chronicity of insomnia, limits the scope of some of our findings and inhibited analysis depth. Additionally, methodological quality was moderate but inconsistent, and insomnia studies were lower quality, highlighting a major limitation of the body of evidence in brain oscillations during sleep. Sufficiently well-powered, sex-stratified analyses are needed to clarify sex effects on NREM oscillations, particularly in insomnia. Greater standardisation of study and brain-oscillation analysis methods would improve the reliability, consistency and comparability of findings across studies. To offer a more comprehensive and sex-sensitive approach, we recommend studies report both absolute and relative power and include event-based measures.

Sex differences are rarely prioritized, often examined in underpowered secondary analyses with unbalanced samples, and group-by-sex interactions are seldom tested. Despite longstanding calls for more sex balanced samples, this continues to be an area of further attention and development in sleep research. Historically, females were excluded from sleep research due to hormonal variability, resulting in a male-biased evidence base [155]. Though both sexes are now included, many studies still overlook hormonal influences which are important for improving our understanding sleep neurophysiology and insomnia risk. This shift from exclusion to unrestricted inclusion may limit progress to better characterize and understand sleep. This can be circumscribed sex-informed, hormonally aware methods, using adequately powered, stratified designs to clarify how sex, age, and insomnia interact to shape sleep neurophysiology.

Our systematic review primarily focuses on biological sex rather than gender (i.e., a societal construct) and gender identity (i.e., a personal construct), all of which can significantly impact sleep[156,157]. Gender-related stressors and discrimination contribute to higher insomnia rates in women and non-cisgender persons, yet neurophysiological studies in these groups are scarce[158]. There is a need for sex inclusive (e.g., how androgens shape sleep dynamics across the lifespan) and gender-inclusive biopsychosocial investigations in sleep science.

## Conclusion

Sex, hormones, and age each play a pivotal role in shaping sleep neurophysiology, yet they remain underexamined in brain-oscillations sleep research, especially in the context of insomnia. Research in this area is increasingly important, as highlighted by Graham and colleagues "Statement on the Importance of Sexuality and Gender Research”[159] released in 2025 which emphasizes the need to understand how these factors intersect with health, behavior, and social well-being. With evolving societal attitudes and growing recognition of gender diversity, it is increasingly essential to expand research in this field. Future research can adopt sex-and gender-informed methods, control for hormonal status, and stratify by age. This can clarify whether true sex-based differences exist in insomnia-related oscillatory activity and support more precise, personalized insomnia interventions that address health disparities. Such work can play a crucial role in advancing health outcomes and ensuring that clinical interventions are inclusive and tailored to the diverse needs of all persons, ultimately, helping to enhance sleep and cognitive outcomes across the lifespan.

### Practice Points

1. Female normal sleepers generally exhibit greater spindle density, absolute sigma power, and slow wave activity (absolute and relative combined) compared to males. There is some but little evidence for amplitude.
2. There is some evidence for normal sleepers having greater spindle density than persons with insomnia, but insomnia subtypes might be a confounder.
3. Presence of heterogeneity in insomnia profiles might influence brain oscillations characteristics and related sex differences
4. Generally, delta power and spindle characteristics (i.e., duration, frequency, amplitude) do not differ between insomnia and normal sleepers.
5. Females and males with insomnia do not differ in either absolute or relative sigma and delta power. However, most studies analysed sex-based differences by combining samples of normal sleepers and persons with insomnia.
6. Few studies investigated sex-by-group interactions in brain oscillations, limiting interpretation.
7. Gonadal hormones appear to impact spindle activity more than slow wave activity, but more research considering hormonal levels and hormonal status are needed.
8. Age, sex, and presence of insomnia impact oscillatory activity.
9. Research on sex-based differences in slow wave and spindle activity remains limited, particularly within insomnia and most is unsatisfactory quality.

## Author Contributions

Conceptualization, N.A.W., A.A.P.; Data curation: N.A.W., E.M.P.; Formal analyses: N.A.W., A.P.; Visualization – A.A.P, A.P., N.A.W.; Supervision: A.A.P., T.D.V.; Writing – original draft preparation, N.A.W., A.A.P.; Writing – review and editing, E.M.P., N.E.C., A.P., T.D.V.; All authors have read and agreed to the published version of the manuscript.

## Other Materials

The data collection forms, extracted data, analytic code, and other materials used in this review are not publicly available. However, all relevant study details and data supporting the findings of this review can be obtained upon request from the corresponding author.

## Acknowledgements

N.A.W received financial support from the Social Sciences and Humanities Research Council and local scholarships (Concordia University) for this project.

## Glossary of terms

**Biological Sex**- the physical and physiological attributes (e.g., chromosomes, hormone levels, reproductive organs, and genitalia) typically categorized as male or female at birth. While most individuals are classified as either male or female, a small percentage are born intersex, meaning they have natural variations in these attributes that do not fit typical biological sex definitions.

Biological sex may not align with an individual’s gender or gender identity.

**Gender-** a societal construct that encompasses roles, behaviors, expectations, and norms culturally associated with being a man or woman. Varies across cultures and time.

**Gender Identity-** a personal, self-defined, internal sense of one’s own gender, which may or may not align with the sex assigned at birth. It includes identities such as man, woman, both, neither, or somewhere along the gender spectrum. May differ from both their biological sex and societal gender expectations.

## Abbreviations

CBTi: Cognitive-Behavioral Therapy for Insomnia
EEG: Electroencephalography
EMG: Electromyography
EOG: Electrooculography
F: Female
NS: Normal sleeper
IN: Insomnia
M: Male
MeSH: Medical Subject Headings
NOS: Newcastle-Ottawa Scale
NREM: Non-Rapid Eye Movement
NREM2: Non-Rapid Eye Movement Stage 2
NREM3: Non-Rapid Eye Movement Stage 3 OA Older Adult
PRISMA: Preferred Reporting Items for Systematic Reviews and Meta-Analyses
PSG: Polysomnography
SO: Slow Oscillation
SW: Slow Wave
SWA: Slow Wave Activity
YA: Young Adult

