## Supplemental Material for "A Systematic Review and Meta-Analysis of Biological Sex Differences in Sleep Spindles and Slow Wave Activity in Adults with and without Insomnia"

**Supplementary Material**

***Figure S1: Search Strategy***

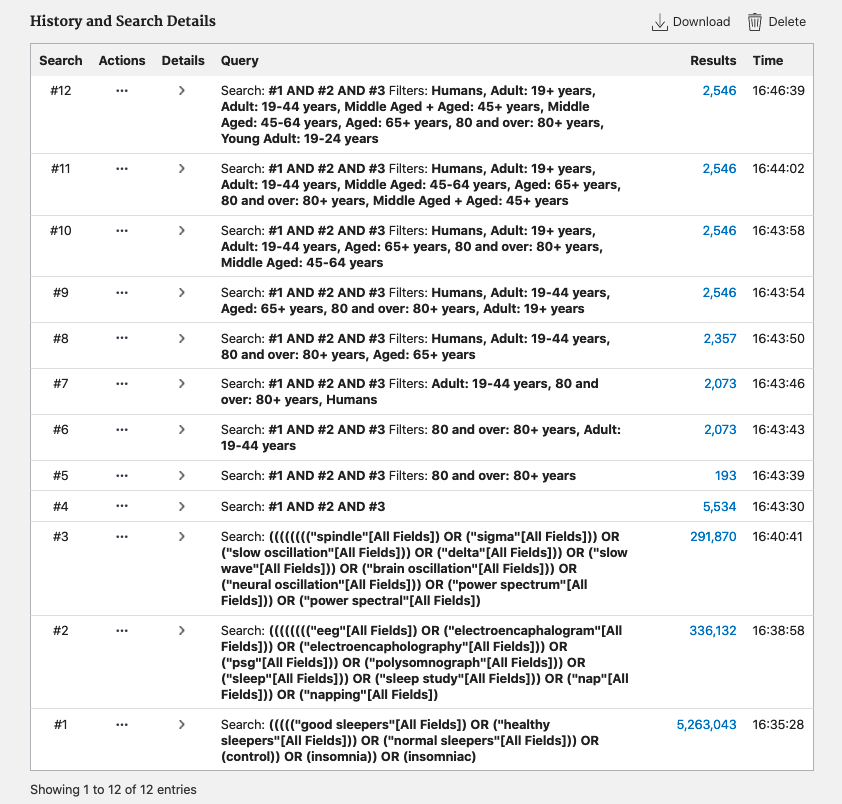

***Underrepresentation of Sex-Based Analyses in Insomnia Research***

In the original search, 32 studies were excluded as they focused only on one sex. Of the studies retained (n= 257), 83% matched the criteria for population and outcomes but did not analyse sex difference (n= 214). Among the eligible studies 14 different countries considered sex differences in brain oscillations in NS and/or individuals with IN. Of the 214 studies that match criteria for population and outcomes but did not analyses sex difference per se, 97 did not list a reason, 64 adjusted for sex (7 articles used sex as a covariate, 8 collected similar sex distribution controls, 47 collected sex-matched controls [2 of which acknowledged the differences by sex in the introduction], 1 paper had all participants in the same menstrual phase, and 1 paper had all females on contraceptives), 5 compared sex for different variables (e.g., %NREM2) but not brain oscillations, and 10 papers acknowledged the impact of sex on brain oscillations in the discussion (7 articles listed not considering sex differences as a limitation, acknowledging the potential impact of female cortisol hormonal fluctuations on brain oscillations [e.g., menstrual cycle phase, menopause, and hormonal contraceptive use], 2 indicated their study was insufficiently powered, and 1 indicated it was outside the scope of the paper).

***Attrition and Acknowledgement of Sex-based Difference per Population Study (NS, IN, NS & IN)***

*Normal Sleepers*

Of the 183 studies on NS that met our eligibility criteria, 33 (18%) directly examined sex-based differences in brain oscillations during sleep, while 150 did not. Of these 150 studies, 56 adjusted for sex in their analyses: 49 used age- and sex-matched controls, 1 further matched participants by menstrual cycle phase, and 6 included sex as a covariate. Among the remaining 94 studies, only 4 provided reasons for not comparing sex-based differences: 2 cited insufficient statistical power, 1 found no significant differences and did not consider sex a confounder, and 1 deemed it outside the study's scope but discussed evidence of sex-based differences in their discussion. The remaining 92 studies did not explain their omission, though 1 noted that gender discrepancies might explain differences in results compared to other studies.

*Insomnia*

Based on our eligibility criteria, 17 studies on individuals with IN were deemed suitable. Of those 17 papers, 2 (12%) directly compared sex-based differences in brain oscillations during sleep, while 15 studies did not consider sex-based differences. One of these 15 studies adjusted for sex in their analyses, although they did not list whether there were sex-based differences in sleep spindles in those with IN. 0/14 of the other studies listed reasons why they did not compare sex-based differences.

*Insomnia and Normal Sleepers*

Lastly, we systematically reviewed papers that contained both NS and those with IN. Based on our eligibility criteria, 41 studies with both NS and those with IN were deemed suitable. Of those 41 papers, 8 (20%) directly compared sex-based differences in brain oscillations during sleep (see Table 3), while 36 studies did not. Of these 36 studies, 16 adjusted for sex in their analyses: 9 used age- and sex-matched controls and 7 included sex as a covariate. Eleven of the 36 studied simply indicated that sex distribution was similar between IN and normal sleeper groups. Among the remaining 9 studies, only 2 provided reasons for not comparing sex-based differences, both citing insufficient statistical power but discussed evidence of sex-based differences in their discussion. The remaining 7 studies did not explain their omission.
